# Symbiosis through lysis: host-triggered prophage induction drives probiotic function in *Lactiplantibacillus plantarum*

**DOI:** 10.64898/2025.12.15.694388

**Authors:** C Zhu, S Manuse, Q Perrier, S Venkatasubramaniyan, H Akherraz, N Lodej, CI Ramos, M Zhang, C Grangeasse, A Razim, F Leulier, RC Matos

**Affiliations:** I Ecole Normale Supérieure de Lyon, Centre National de la Recherche Scientifique, Institut de Génomique Fonctionnelle de Lyon UMR5242 69364 Lyon Cedex 07 France; Molecular Microbiology and Structural Biochemistry, CNRS UMR 5086, Université Claude Bernard Lyon 1, Lyon, France; Centre National de la Recherche Scientifique, Ecole Normale Supérieure de Lyon, Institut de Génomique Fonctionnelle de Lyon UMR5242 69364 Lyon Cedex 07 France; Université Claude Bernard Lyon 1, Ecole Normale Supérieure de Lyon, Centre National de la Recherche Scientifique, Institut de Génomique Fonctionnelle de Lyon UMR5242 69364 Lyon Cedex 07 France; Laboratory of Aquaculture Nutrition and Environmental Health, School of Life Sciences, East China Normal University, Shanghai 200241, China; Hirszfeld Institute of Immunology and Experimental Therapy. Polish Academy of Sciences, Wrocław, Poland

## Abstract

The release of bacterial bioactive molecules across the gut barrier is a crucial yet poorly understood step in microbe-host molecular dialog. Here we unravel a phage-driven mechanism that enables this process in a symbiont that supports host adaptive growth. In *Lactiplantibacillus plantarum* NC8 (*Lp^NC8^*), we identify a stress-inducible prophage, pp2, that undergoes genotoxic-stress-dependent activation *in vivo* and triggers holin-lysin-mediated lysis. This controlled lytic program produces phage particles together with extracellular vesicles (EVs), enriched in symbiotic cues, including lipoteichoic acids. In a model of beneficial symbiosis, pp2-dependent lysis is strictly required for the ability of *Lp^NC8^*to support juvenile growth in nutritionally challenged *Drosophila melanogaster*. Disruption of pp2-dependent lysis abolishes EVs release *in vitro* and alters the growth-promoting effect *in vivo*. We show that the acidic region of the *Drosophila* midgut, but not host antimicrobial peptides nor lysozymes, acts as a physiological trigger of prophage induction: removal of this acidic compartment markedly reduces phage production and impairs *Lp^NC8^*-mediated growth promotion. These findings demonstrate that host gut physiology regulates prophage activation and reveal prophage-induced lysis as a previously unrecognized mechanism by which a beneficial gut bacterium releases symbiotic cues to support host development during nutritional stress.

## INTRODUCTION

Metazoans rely on their resident microorganisms for a wide range of physiological functions, from nutrient acquisition to immune maturation and metabolic homeostasis^1^. During postnatal development, these host-microbe interactions are especially critical: juvenile growth, a period of rapid somatic expansion and organ maturation, requires both adequate nutrition and a properly assembled gut microbiota^2^. Undernutrition perturbs microbial community maturation and leads to impaired linear and ponderal growth^3,4^. Microbial (probiotic) or microbiota-directed nutritional (prebiotic) interventions can mitigate these defects^5^, yet the mechanisms by which gut bacteria contribute to healthy growth and buffer the physiological consequences of undernutrition remain incompletely understood.

*Drosophila melanogaster* provides a powerful model for dissecting the mechanisms and physiological consequences of host-commensal interactions^6,7^. Commensal bacteria associated with *Drosophila*, including members of the genus *Lactiplantibacillus*, modulate multiple aspects of host physiology, notably juvenile growth. Work over the past decade has identified several molecular symbiotic cues, bacterial components that directly modulate host physiology, produced by *L. plantarum*. These include peptidoglycan (PG) fragments sensed by enterocyte PG recognition proteins (PGRPs), which activate the IMD/NF-κB pathway to induce intestinal peptidases and enhance protein assimilation^8^. The *dltEXABCD* operon, which decorates lipoteichoic acids with D-alanine (d-Ala-LTAs), is similarly essential for growth promotion^9,10^. More recently, *L. plantarum* ribosomal and transfer RNAs (r/tRNAs) were shown to activate the host General Control Nonderepressible 2 kinase (GCN2) in a subset of larval enterocytes, remodelling the intestinal transcriptome to support systemic anabolic growth^11^. Importantly, cell envelope components of *L. plantarum*, including PG, are sufficient to promote growth in undernourished mice, suggesting that these molecular dialogs reflect an evolutionarily conserved mechanism of microbe-mediated juvenile growth^12^.

Despite this progress, a key mechanistic question remains unresolved: How do these diverse symbiotic cues, PG fragments, d-Ala-LTAs, and r/tRNAs, reach host intestinal cells? Most are either intracellular or tightly associated with the bacterial cell envelope, and the architecture of the gut generally restricts direct contact between bacteria and the epithelium^13,14^. No dedicated secretion system is known in *L. plantarum* that could export these molecules in physiologically relevant amounts. This gap suggests that release into the intestinal lumen may depend on mechanisms of bacterial lysis. Bacterial lysis can occur through multiple processes, but prophage induction represents a conserved, environmentally responsive route. Prophages, integrated viral genomes, remain transcriptionally silent during lysogeny but can be induced by stressors such as DNA damage, nutrient limitation, or activation of the SOS response. Upon induction, prophages initiate their lytic program, culminating in bacterial cell lysis and the release of phage particles together with intracellular contents and cell envelope fragments that can also form extracellular vesicles^15^. In the gut environment, where bacteria are exposed to physicochemical and immune constraints, prophage induction may therefore constitute a key mechanism linking host intestinal physiology to bacterial secretion dynamics^16–18^.

The *Drosophila* gut provides several potential triggers for such processes. The larval midgut contains a highly acidic compartment, the copper cell region, which functionally resembles the mammalian stomach and exerts strong bactericidal activity. In parallel, intestinal epithelial cells produce antimicrobial peptides and lysozymes that respectively regulate microbial colonization and/or remodel bacterial cell walls^19^. Together, these host-derived processes create an intestinal environment that may promote prophage induction, bacterial cell lysis, and vesicle release in commensal bacteria. Yet whether prophage-mediated lysis benefits host physiology, particularly by enabling the delivery of symbiotic cues, remains largely unexplored.

Here, we identify a single inducible prophage, pp2, in the beneficial *L. plantarum* NC8 strain as a necessary and sufficient driver of environmentally responsive bacterial lysis, extracellular vesicle release and microbe-mediated host growth. We further show that the acidic region of the *Drosophila* gut influences prophage induction and phage release *in vivo*, directly linking host intestinal physiology to bacterial lysis dynamics and ultimately host benefits. Our findings reveal prophage-mediated lysis as a mechanism through which a beneficial *L. plantarum* strain delivers symbiotic cues to the host and benefits its physiology, uncovering phage-driven lysis as a previously unrecognized determinant of probiotic function.

## RESULTS

### Prophages are important for *Lp* probiotic properties

To explore the prophage landscape of *Lactiplantibacillus plantarum* NC8 (*Lp^NC8^*), we first interrogated its genome using PHASTEST, a computational tool that identifies and annotates prophage sequences within bacterial chromosomes^20^. Temperate bacteriophages typically exhibit a modular genomic architecture in which clusters of genes encode distinct functional units corresponding to key stages of the phage life cycle. Classical analyses recognize six of such modules - lysogeny, replication, transcriptional regulation, head and tail morphogenesis, DNA packaging, and cell lysis - which together impose a temporal order on phage gene expression^21^. PHASTEST identified three prophage regions within the *Lp^NC8^* genome: prophage 1 (pp1; 16.9 kb), prophage 2 (pp2; 41.8 kb), and prophage 3 (pp3; 43.4 kb) (Fig. 1a and Supplementary Fig. 1). Based on their gene content, pp2 and pp3 are predicted to be intact prophages, as each contains the full set of functional modules characteristic of complete temperate phage genomes. In contrast, pp1 exhibits a reduced architecture: although it encodes an integrase, replication protein, head morphogenesis components, and DNA-packaging machinery, it lacks genes required for tail morphogenesis and host-cell lysis. This organization mirrors that of a recently described family of phage-inducible chromosomal islands (PICIs) termed capsid-forming PICIs (cf-PICIs)^22^. cf-PICIs retain the genetic determinants necessary for assembling small capsids and for selectively packaging their own genomes, but they depend on helper phages to supply tail structures required for completing infectious particles^22^.

**Fig. 1.**
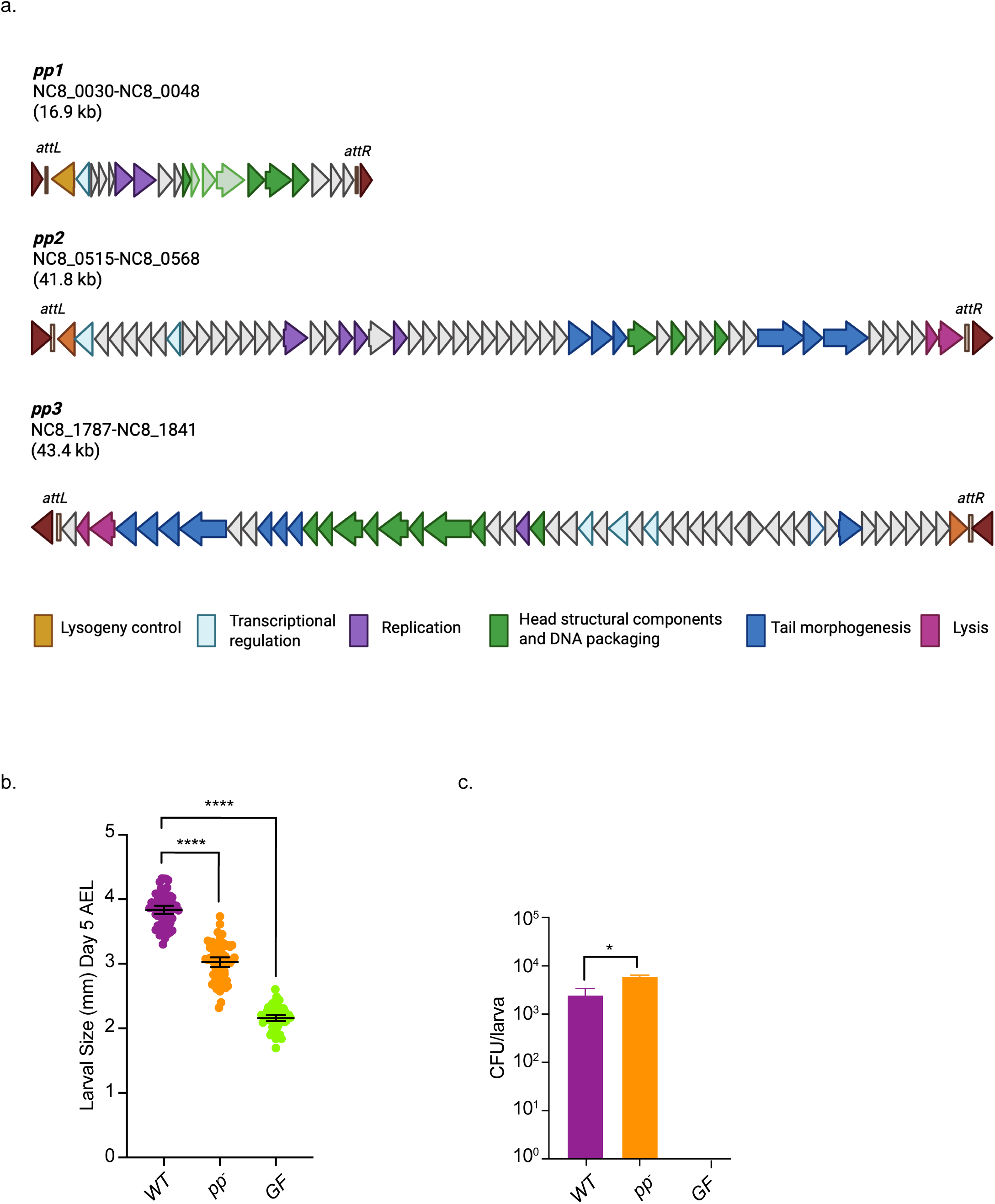
Prophages are important to *Lp^NC8^* probiotic traits. (a) *Lp^NC8^*prophages as predicted by PHASTEST. Open reading frames are indicated by arrows. Only genes encoding predicted functions are annotated. Colors correspond to the functional models of temperate phages as depicted on the bottom of the figure. (b) Larval longitudinal length of eggs associated with *Lp^NC8^*(n=60), *pp^-^* (n=60) strains or PBS (n=60) (GF), 5 days after egg laying (AEL) on low-protein fly food (7g/L of yeast). Asterisks represent statistically significant difference compared to *Lp^NC8^* (WT) larval size. Significance was determined using Kruskal-Wallis. ****P<0.0001. The bars in the graph represent mean and 95% CI. A representative graph from one out of three independent experiments is shown. (c) Bacterial load of *Lp^NC8^*(n=3) and *pp^-^* (n=3) strains recovered from larvae associated with 10^7^ CFUs after 5 days association. Asterisks represent statistically significant difference compared to *Lp^NC8^* (WT) CFUs. *P<0.05. The bars in the graph represent mean and 95% CI. A representative graph from one out of three independent experiments is shown.

To assess the global contribution of *Lp^NC8^* prophages to host physiology, we generated a prophage-free strain (*pp-*) in which all three prophage regions were deleted, resulting in a cumulative 102-kb chromosomal deletion (Supplementary Fig. 1). We then compared the probiotic activity of the wild-type strain and the *pp-* mutant in a *Drosophila* undernutrition model, focusing on their ability to promote host juvenile growth despite undernutrition (Fig. 1b). Quantification of larval size at day five after bacterial association, demonstrated that removal of prophages markedly reduced the growth-promoting activity of *Lp^NC8^*. Importantly, both strains colonized the fly diet to similar levels, even slightly improved for the *pp-* mutant (Fig. 1c), indicating that the reduced growth promotion of the *pp-* strain is not attributable to impaired persistence. These findings suggest that prophages contribute to the probiotic capacity of *Lp^NC8^*, prompting us to investigate their mechanistic role in host growth promotion.

### *Lp^NC8^* prophages induction is triggered upon genotoxic stress

To assess the activity of *Lp^NC8^* prophages and their capacity to form functional phage particles, exponentially growing cultures were treated with mitomycin C, a DNA-damaging agent that activates the bacterial SOS response and thereby induces prophages to enter the lytic cycle. The SOS response is a highly conserved bacterial pathway that orchestrates the expression of genes involved in DNA repair and stress adaptation, primarily through the regulatory activities of LexA and RecA^23^. Under lysogenic conditions, phage lytic genes are repressed by LexA, maintaining prophages in a dormant state. In the absence of RecA, the LexA co-protease is not activated, impairing the initiation of the lytic program and consequently reducing prophage induction and host-cell lysis^16^. To detect functional prophage activity, we employed the prophage-free strain as an indicator (sensitive bacterial host on which phages can form plaques), enabling plaque formation by prophages capable of excision, replication, and virion production. In addition to wild-type *Lp^NC8^* and the prophage-deleted strain (*pp-),* we included a Δ*recA* mutant, which is deficient in SOS-mediated induction, to assess the dependence of prophage activation on RecA function. Mitomycin C treatment of *Lp^NC8^* resulted in a ∼5-log reduction in colony-forming units (CFUs), accompanied by an increase in plaque-forming units (PFUs) (Fig. 2a). Importantly, we also detected PFUs at steady state revealing basal phage lytic activity. These observations indicate that *Lp^NC8^* cells undergo lysis, releasing phage particles capable of forming plaques on the prophage-free indicator strain. By contrast, the *pp-* strain produced no active phage particles despite a ∼3-log decline in CFUs. Similarly, the Δ*recA* mutant was unable to activate the SOS response, preventing prophage derepression, lytic induction, and phage production, as reflected by the absence of plaques (Fig. 2a).

**Fig. 2.**
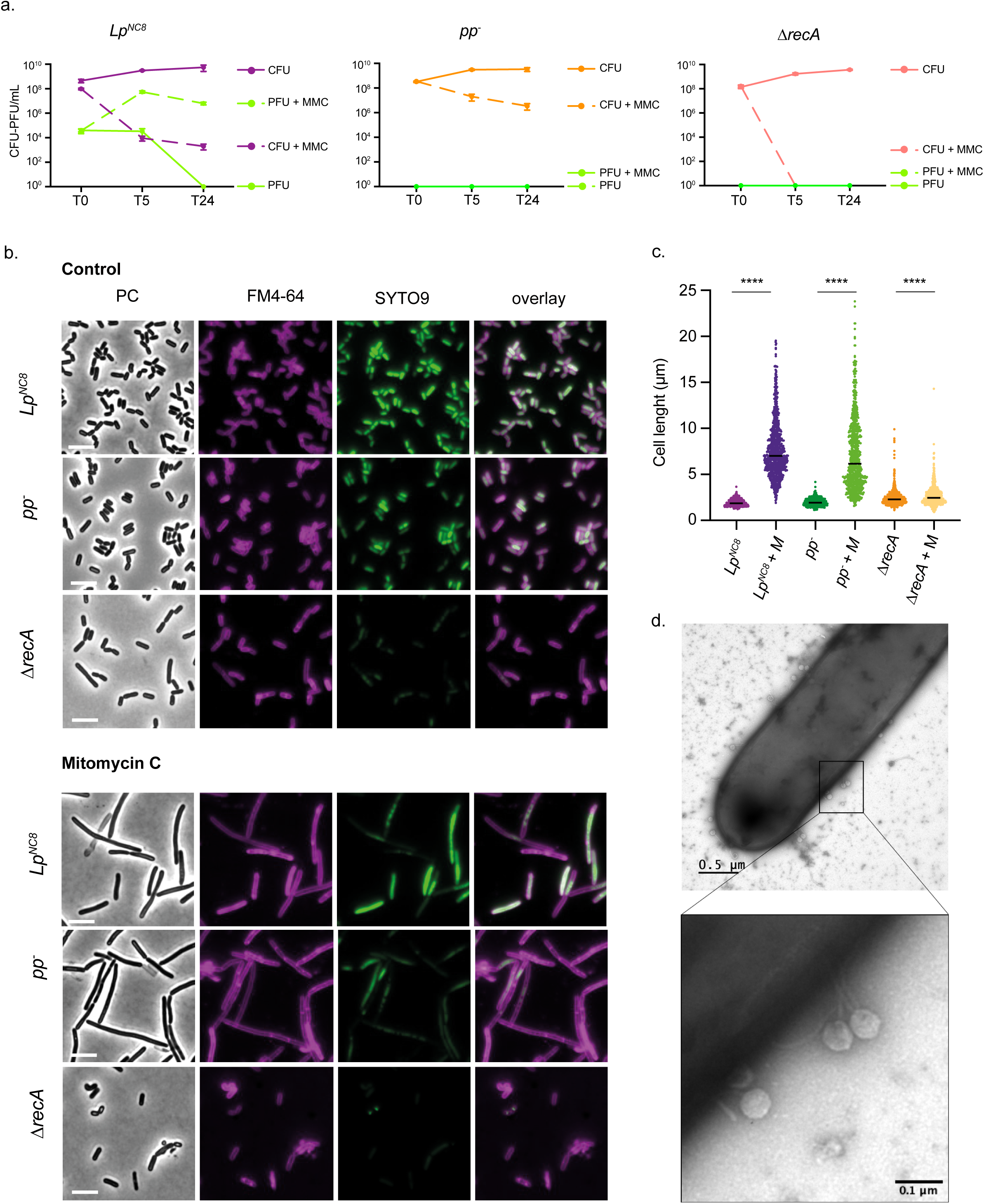
*Lp^NC8^* prophages induction is increased upon genotoxic stress. (a) CFUs *vs* PFU representation for *Lp^NC8^*, *pp*^-^ and *ΔrecA* strains. Exponentially growing cultures were induced with mitomycin C. T0 corresponds to the time of mitomycin C supplementation. Cells and supernatants were harvested at different timepoints after induction and analyzed for CFUs and PFUs on a lawn of *pp^-^* strain, respectively. (b) Fluorescent microscopy: representative images of *Lp* cultures (*Lp^NC8^*, *pp^-^, ΔrecA*) treated or not with mitomycin C, and dyed with FM4-64 (false-colored in magenta) to stain membranes. PC stands for phase contrast. Scale bars, 5 µm. (c) Cell length: quantification of cell length per single cell across *Lp* cultures (*Lp^NC8^*, *pp^-^*, *ΔrecA*) treated or not with mitomycin C. For each sample, *n* = 800 cells were randomly selected from larger dataset. Thick lanes represent the median. Significance was determined using Kruskal-Wallis and Dunn’s multiple comparison tests. ****P<0.0001. (d) Transmission electron microscopy of negative stained *Lp^NC8^* cells after mitomycin C supplementation.

To further assess cell viability following mitomycin C-induced prophage activation, we employed the LIVE/DEAD® BacLight™ Bacterial Viability Kit to quantify the proportion of non-viable cells across the different strains (Supplementary Fig. 2). This complementary analysis corroborates the CFU and PFU data shown in Fig. 2a, revealing that approximately 40% of *Lp^NC8^*cells are non-viable after mitomycin C treatment. In contrast, only ∼10% of cells in the prophage-free strain succumb under the same conditions. To determine whether prophage induction is enhanced by other genotoxic stressors, we exposed cultures to other DNA-damaging agents, including ciprofloxacin and colibactin (produced by *E. coli* in co-culture with *Lp^NC8^* ^18^), as well as to the non-genotoxic antibiotic polymyxin B. Both ciprofloxacin and colibactin markedly increased PFUs production while reducing CFUs, consistent with robust prophage induction. In contrast, polymyxin B did not alter phage output, which remained at spontaneous-induction levels (Supplementary Fig. 3). These experiments indicate that genotoxic stress is a driver of prophage induction *in vitro*.

We next used fluorescence microscopy to examine whether prophage induction by mitomycin C affects cell morphology. Cells were stained with FM4-64, a lipophilic dye that labels cell membranes. As shown in Fig. 2b-c, mitomycin C treatment induced pronounced filamentation in both wild-type *Lp^NC8^*and *pp-* strains, producing cells that were substantially longer than non-induced controls. This filamentous phenotype was absent in the Δ*recA* mutant, indicating that cell elongation is dependent on SOS response activation; in the absence of RecA, the SOS regulon cannot be induced. Collectively, these observations demonstrate that genotoxic stress alters cellular morphology and coincides with increased production of infectious phage particles. Finally, we visualized *Lp^NC8^* supernatants following mitomycin C induction by transmission electron microscopy with negative staining (Fig. 2d). Phage particles were observed being released from lysed cells, providing direct evidence that *Lp^NC8^* produces active virions upon prophage induction. Together, these findings demonstrate that *Lp^NC8^* prophages are inducible under genotoxic stress, reshaping cellular morphology and driving the production of infectious phage particles.

### Identification of pp2 as the active prophage driving virion production in *Lp^NC8^*

As noted above, only prophages pp2 and pp3 are predicted to be fully functional based on PHASTEST analysis (Fig. 1a). Having established that *Lp^NC8^* carries inducible prophages capable of entering the lytic cycle and producing infectious virions, we next sought to determine which specific prophages are active. To address this, we generated a panel of isogenic strains designed to allow assessment of individual phage activity through plaque assays using tailored indicator strains (Supplementary Fig. 1). We first engineered monolysogenic *Lp^NC8^* strains, each retaining only a single prophage - *pp1⁺*, *pp2⁺*, or *pp3⁺* - and assessed their activity using plaque assay on the prophage-free (*pp-*) indicator strain. Supernatants from mitomycin C-induced cultures of each monolysogen were spotted onto *pp-* lawns. Plaques were observed exclusively with the *pp2⁺* supernatant, demonstrating that pp2 is the sole prophage capable of producing infectious particles (Fig. 3a).

**Fig. 3.**
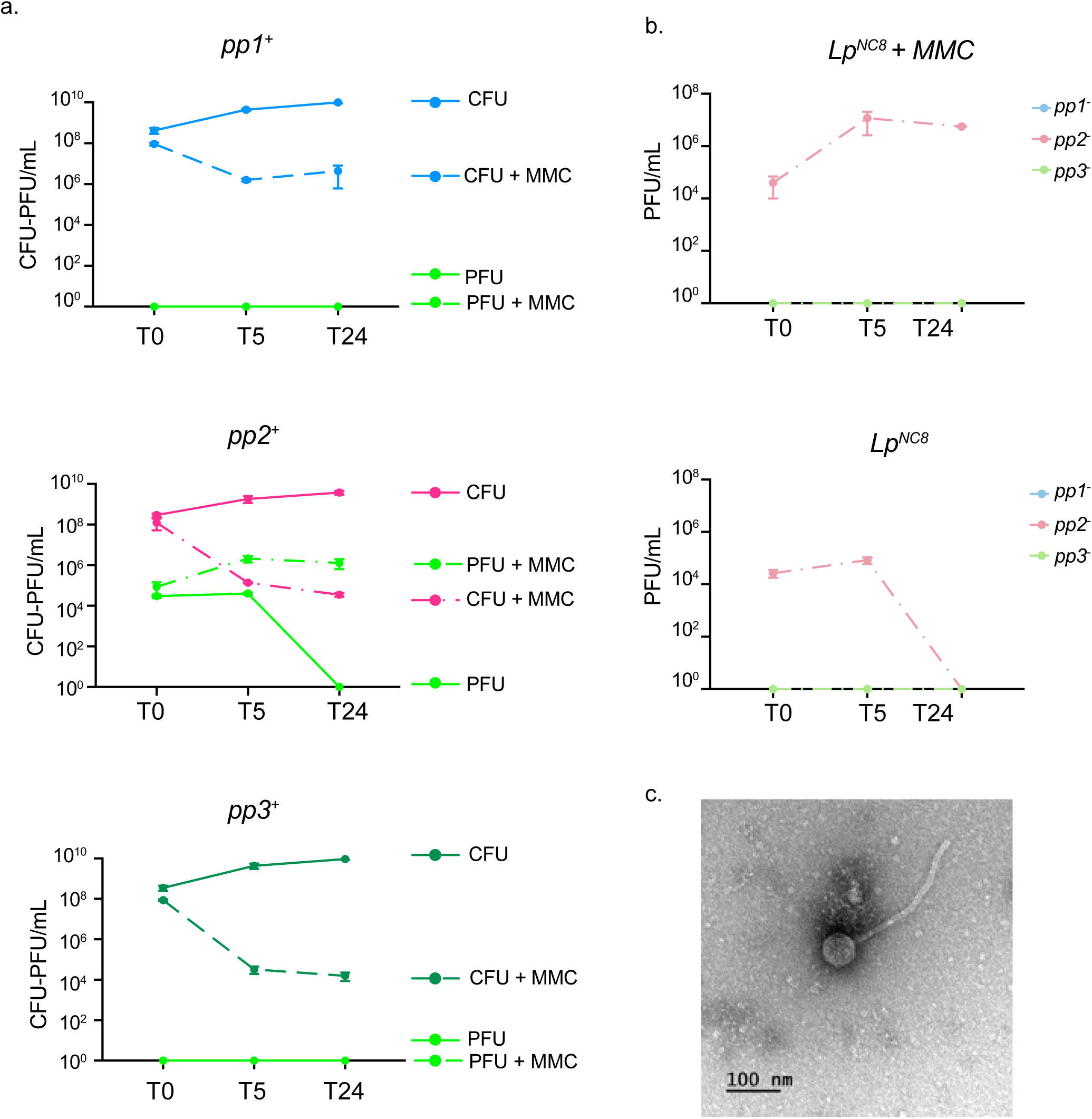
Prophage 2 forms infectious particles. (a) CFUs *vs* PFU representation for monolysogenic strains, *pp1^+^*, *pp2^+^*, *pp3^+^*, upon mitomycin C supplementation. Exponentially growing cultures were induced with mitomycin C. T0 corresponds to the time of mitomycin C supplementation. Cells and supernatants were harvested at different timepoints after induction and analyzed for CFUs and PFUs on a lawn of *pp^-^*strain, respectively. (b) Superinfection assay: PFUs representation from *Lp^NC8^*supernatant on *pp1^-^*, *pp2^-^* and *pp3^-^*strains lawn, with and without mitomycin C. Exponentially growing *Lp^NC8^*cultures were induced with mitomycin C. T0 corresponds to the time of mitomycin C supplementation. Supernatants were harvested at different timepoints after induction and analyzed for CFUs and PFUs on a lawn of *pp1^-^*, *pp2^-^* and *pp3^-^* strains, respectively. (c) *pp2^+^*supernatant harvested upon mitomycin C induction observed by TEM with negative staining.

To further confirm that pp2 is the only prophage capable of lytic induction and virion production, we performed targeted superinfection assays using *Lp^NC8^* strains each lacking a single prophage (*pp1-*, *pp2-*, or *pp3-*). In this setup, each deletion strain should be resistant to all prophages present in the wild-type genome except the one it lacks. When supernatants from mitomycin C-induced wild-type *Lp^NC8^*were applied, plaques formed exclusively on the *pp2^-^* strain, demonstrating that only pp2 generates infectious particles (Fig. 3b). Transmission electron microscopy of negatively stained *pp2⁺* supernatants revealed virions with the characteristic morphology of the *Siphoviridae* family, including an icosahedral head and a long, non-contractile flexible tail^24^ (Fig. 3c). These results provide compelling evidence for pp2 as the sole active prophage within *Lp^NC8^* genome.

### pp2-induced lysis drives formation of extracellular vesicles in *Lp^NC8^*

Our data indicate that pp2 is the only *Lp^NC8^* prophage capable of producing infectious phage particles and, accordingly, is predicted to lyse the host cell to complete a full lytic cycle. Prophage-mediated lysis is generally executed by a holin-lysin module encoded within the prophage genome^25^. To test whether pp2 is required for *Lp^NC8^* cell lysis, we deleted the holin-lysin genes from pp2 in both the wild-type *Lp^NC8^* background (*Lp^NC8^_Δpp2lysis_*) and the pp2 monolysogenic strain (*pp2^+^_Δpp2lysis_*). Given that lysis is a tightly regulated, temporally coordinated process that ensures timely release of phage progeny^25^, we anticipated that removal of the lysis module would impair both cell lysis and the production of infectious particles. We next induced exponentially growing cultures of the lysis-deficient and control strains and quantified CFUs and PFUs (Fig. 4a). Consistent with previous observations, wild-type *Lp^NC8^* and the *pp2^+^*monolysogen exhibited a decline in CFUs accompanied by an increase in PFUs upon mitomycin C treatment, confirming that pp2 induction triggers cell lysis and phage release (Fig. 1 and 2). In contrast, deletion of the pp2 holin-lysin module in either the *Lp^NC8^* or *pp2^+^* monolysogen backgrounds completely abolished PFU formation following mitomycin C induction (Fig. 4a). These results confirm that pp2 and its lysis module are essential for executing cell lysis and releasing infectious phage particles. Surprisingly, deletion of the pp2 holin-lysin module did not enhance bacterial survival. Although unexpected, this phenotype may reflect the accumulation of phage particles retained within the cells, which could impose an additional physiological burden and contribute to cell death despite the absence of lytic release.

**Fig. 4.**
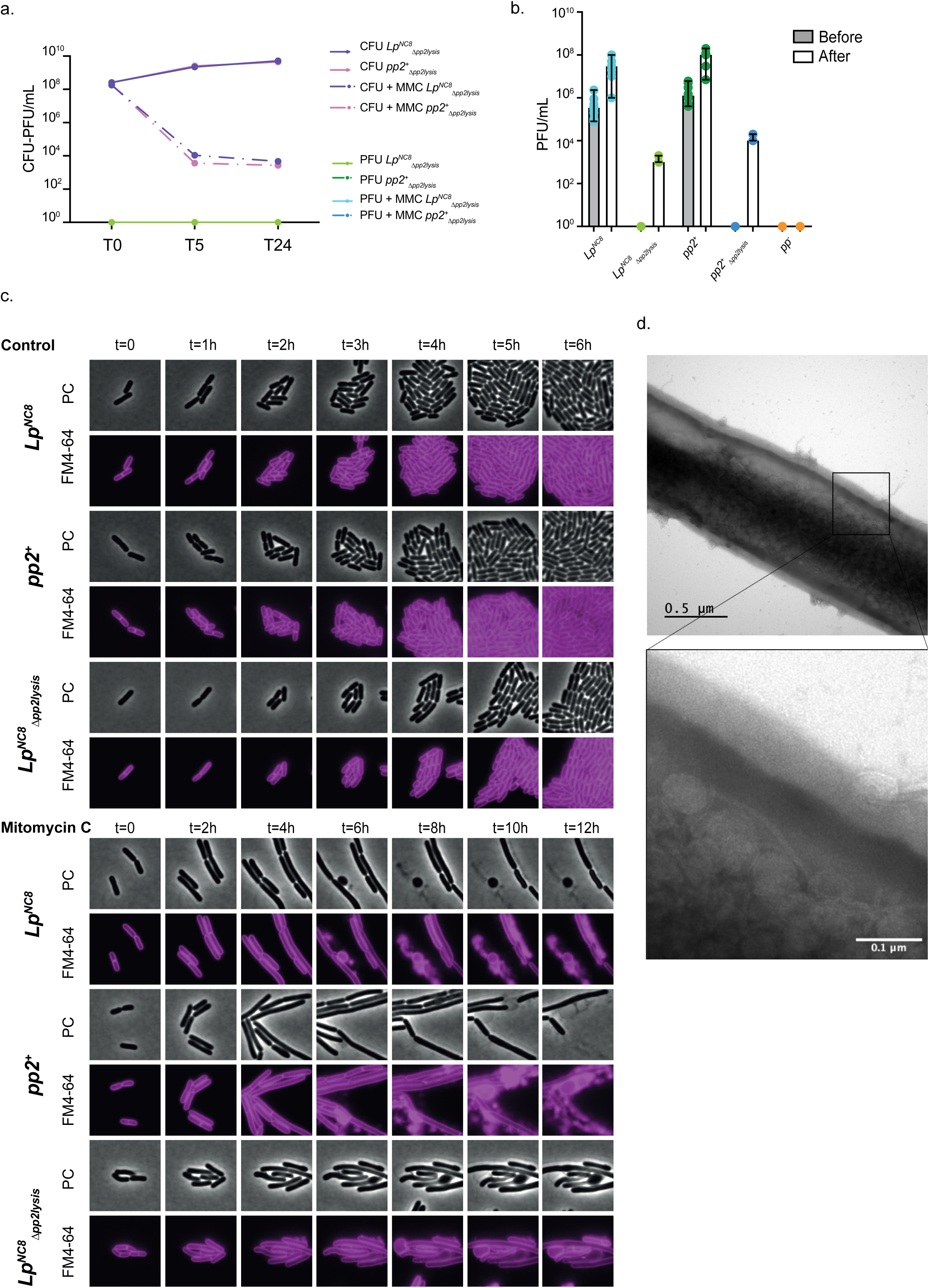
Genotoxic stress triggers pp2-dependent cell lysis and particle formation. (a) CFUs *vs* PFU representation of *Lp^NC8^*, *pp2^+^*strains. Exponentially growing cultures were induced with mitomycin C. T0 corresponds to the time of mitomycin C supplementation. Cells and supernatants were harvested at different timepoints after induction and analyzed for CFUs and PFUs on a lawn of *pp^-^* strain, respectively; (b) Lysozyme digestion assay performed on wild-type *Lp^NC8^*, the pp2 monolysogen (*pp2^+^*), the prophage-free strain (*pp^-^*), and their corresponding lysis-deficient derivatives (*Lp^NC8^* and *pp2^+^*). Cultures were subjected or not to exogenous lysozyme treatment (5 mg.mL^-1^, 1 h), after which released phage particles in the supernatant were quantified by plaque assay. PFUs were obtained from 3 independent experiments. The bars in the graph represent mean and 95% CI. (c) Time lapse of *Lp^NC8^*, *pp2^+^*, *Lp^NC8^*strains. Time lapse montages of selected fields of view from Movies ABC, showing bacterial shape alterations under phage induction. Cells were placed on agarose pads containing the fluorescent membrane dye FM4-64 (false-colored in magenta), with or without mitomycin C, and observed at 30°C. Scale bars, 5 µm. (d) Transmission electron microscopy image of negative stained *Lp^NC8^*cells.

To test this hypothesis, we artificially lysed bacterial cells using the peptidoglycan-digesting enzyme lysozyme (Fig. 4b). We first quantified PFUs in the supernatants of mitomycin C-induced cultures prior to lysozyme treatment. After collecting these initial supernatants, the remaining cells were exposed to lysozyme, and PFUs were enumerated again. This strategy allowed us to distinguish between phages released via prophage-induced lysis and those retained intracellularly, which were liberated following enzymatic digestion of the cell wall. As expected, supernatants from mitomycin C-treated wild-type *Lp^NC8^* and *pp2^+^* strains contained phage particles capable of forming plaques on the indicator strain. Remarkably, lysozyme treatment of strains lacking the pp2 lysis module also resulted in PFU formation, demonstrating that phages were indeed produced but remained trapped inside the host cells. These results provide further evidence that the pp2 holin-lysin module is essential for efficient bacterial lysis and the release of infectious particles in *Lp^NC8^*.

Building on the critical role of the pp2 holin-lysin module in *Lp^NC8^*cell lysis, and considering recent reports connecting phage-induced lysis to extracellular vesicles release in Gram-positive bacteria^26,27^, we investigated whether pp2-mediated lysis also drives the formation of extracellular vesicles in *Lp^NC8^*. To address this, we monitored the growth kinetics of *Lp^NC8^* strains with or without the pp2 lysis module (*Lp^NC8^*, *pp2^+^*, *Lp^NC8^_Δpp2lysis_*) over 12 h in the presence or absence of mitomycin C, and labeled membranes with the membrane lipophilic dye FM4-64, which stains both cells and potential particles. Under normal growth conditions (MRS supplemented with FM4-64), all strains divided approximately every hour and reached stationary phase without detectable lysis or extracellular particle formation (Fig. 4c; Supplementary Video 1-3). Strikingly, in the presence of mitomycin C, wild-type *Lp^NC8^* and the *pp2^+^* monolysogen ceased division and elongated. Approximately six hours post-induction, these cells underwent lysis, coinciding with the appearance of round, membrane-bound structures resembling bacterial extracellular vesicles (Fig. 4c; Supplementary Video 4 and 5). The observation of this phenotype in the *pp2^+^* monolysogen confirms that pp2 alone drives both cell lysis and particle formation. By contrast, deletion of the *pp2* lysis module in *Lp^NC8^* (*Lp^NC8^_Δpp2lysis_*) impaired bacterial lysis and formation of extracellular particles (Fig. 4c; Supplementary Video 6). Notably, few cells displayed a distinctive spoon-shaped morphology (Supplementary Fig. 4), potentially reflecting intracellular accumulation of phage particles concentrated at one cell pole. This interpretation is consistent with our previous finding that enzymatic digestion of the cell wall restores phage release in lysis-deficient strains (Fig. 4b). Consistently, transmission electron microscopy (TEM) of *Lp^NC8^* cells reveals phage particles retained within intact, non-lysing cells (Fig. 4d).

To determine the nature of the observed extracellular particles, we isolated, purified, and characterized samples following the MISEV2023 guidelines^28^. We combined ultracentrifugation with size-exclusion chromatography (SEC), TEM, particle sizing, and LTA enrichment, an approach commonly used to operationally define Gram-positive membrane-derived vesicles^29^. TEM analysis of mitomycin-induced *Lp^NC8^*cultures revealed abundant phage heads, tails, and additional extracellular particles. Notably, these non-phage particles were also observed in non-induced *Lp^NC8^* samples as well as in the prophage-free *pp^−^*strain lacking all prophage genes (Fig. 5a). Particle quantification analysis revealed a significant increase in particle abundance following mitomycin C treatment, while only minor differences in mean particle size were observed between conditions (Fig. 5b-c). Importantly, the presence of prophages in *Lp^NC8^* was associated with a higher number of released vesicles compared to the *pp^−^*strain, indicating that phage-mediated lysis strongly enhances vesicle production. However, mitomycin C treatment also increased vesicle release in the *pp^−^* background, suggesting that additional mechanisms independent of prophages activation contribute to vesiculogenesis. The concomitant decrease in CFU counts observed after induction suggests that a substantial proportion of these particles originate from bacterial cell lysis (Fig. 5d). Finally, analysis of lipoteichoic acids (LTAs), a hallmark component of Gram-positive bacterial membranes, confirmed the presence of LTAs in samples of mitomycin C induced *Lp^NC8^* cultures (Fig. 5e). Together, these results support the conclusion that prophage induction promotes the release of extracellular vesicles enriched in bacterial membrane-associated components.

**Fig. 5.**
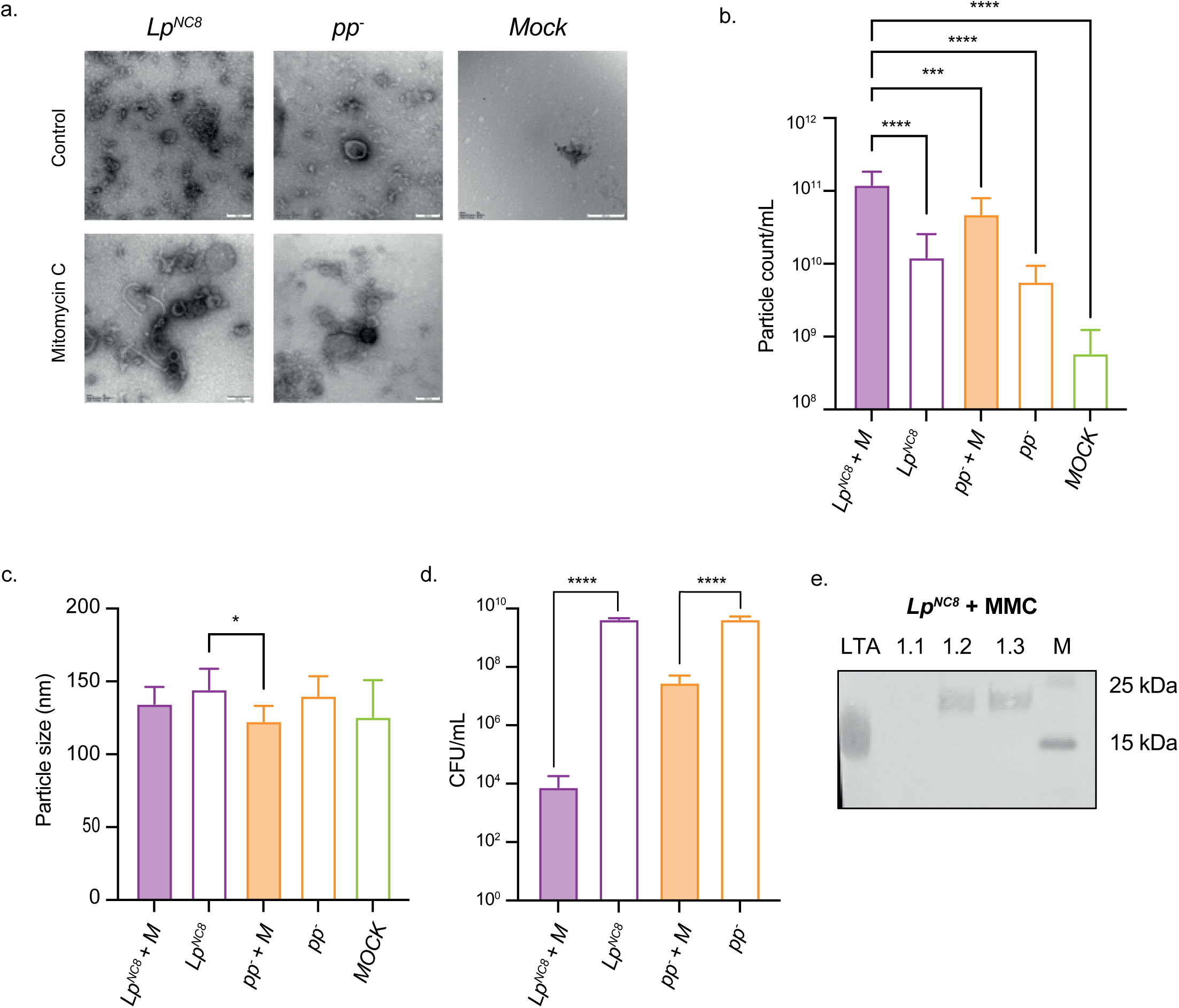
Particles formed by prophage-induced cell lysis are bona-fide bacterial extracellular vesicles. (a) TEM visualization of selected samples from *Lp^NC8^*and *pp^-^* cultures induced and non-induced by mitomycin C. (b) Zetasizer analysis of particle concentration, data shown for three independent biological repeats. Significance was determined using ordinary one-way ANOVA and Tukey’s multiple comparison test. Asterisks illustrate statistically significant differences (p<0.05). (c) Zetasizer analysis of particle size, data shown for three independent biological repeats. Significance was determined using ordinary one-way ANOVA and Tukey’s multiple comparison test. Asterisks illustrate statistically significant differences (p<0.05). (d) CFUs from which vesicles were isolated. Data shown for three independent biological repeats. Significance was determined using ordinary one-way ANOVA and Tukey’s multiple comparison test. Asterisks illustrate statistically significant differences (p<0.05). (e) Western blot analysis of LTA content in analyzed samples. 15 µl of each Size Exclusion Chromatography (SEC) fractions were loaded per well, 25 µg of commercial LTA was used as positive control.

Collectively, these results support a model in which pp2 induction triggers entry into the lytic cycle, production of infectious pp2 virions, and subsequent prophage-mediated cell lysis, during which bacterial extracellular vesicles are formed.

### pp2-induced lysis is required for *Lp^NC8^*-mediated larval growth promotion

Having established that pp2-mediated lysis drives the release of phage and extracellular particles, we next asked whether prophage-induced lysis contributes to the growth-promoting effect of *Lp^NC8^* in undernourished *Drosophila* larvae. As shown in Figure 1b, prophages are required for *Lp^NC8^*-mediated growth promotion. To investigate whether inducible prophage activity and the resulting bacterial lysis mediate the release of symbiotic cues important for supporting host growth, we employed a gnotobiotic *Drosophila* model. Germ-free eggs were inoculated with 10^7^ CFUs of *Lp^NC8^* and *pp2*^+^ (both capable of inducing cell lysis and extracellular particle formation), *Lp^NC8^* and *pp2*⁺ (lysis-deficient strains), or *pp^-^* (lacking all prophages). Larvae were raised on poor diet and collected on days 4, 5, and 6 post-inoculation to measure body size (Fig. 6a), *in vivo* bacterial load (CFUs; Fig. 5b), and *in vivo* phage release (PFUs; Fig. 6c). Larval size measurements revealed that *Lp^NC8^* and pp2⁺ robustly promoted juvenile growth on poor diet, with larvae significantly longer than germ-free controls. By contrast, larvae associated with *Lp^NC8^*, *pp2*⁺ or pp- were markedly smaller. Bacterial loads remained comparable across all conditions (Fig. 6b), indicating that differences in growth promotion were not attributable to altered bacterial colonization of the host. PFU quantification confirmed *in vivo* phage release exclusively in *Lp^NC8^* and *pp2*^+^-associated larvae, increasing between days 5 and 6 and coinciding with maximal host growth promotion (Fig. 6c). Lysis-deficient strains failed to release phages, confirming impaired prophage-mediated lysis *in vivo*.

**Fig. 6.**
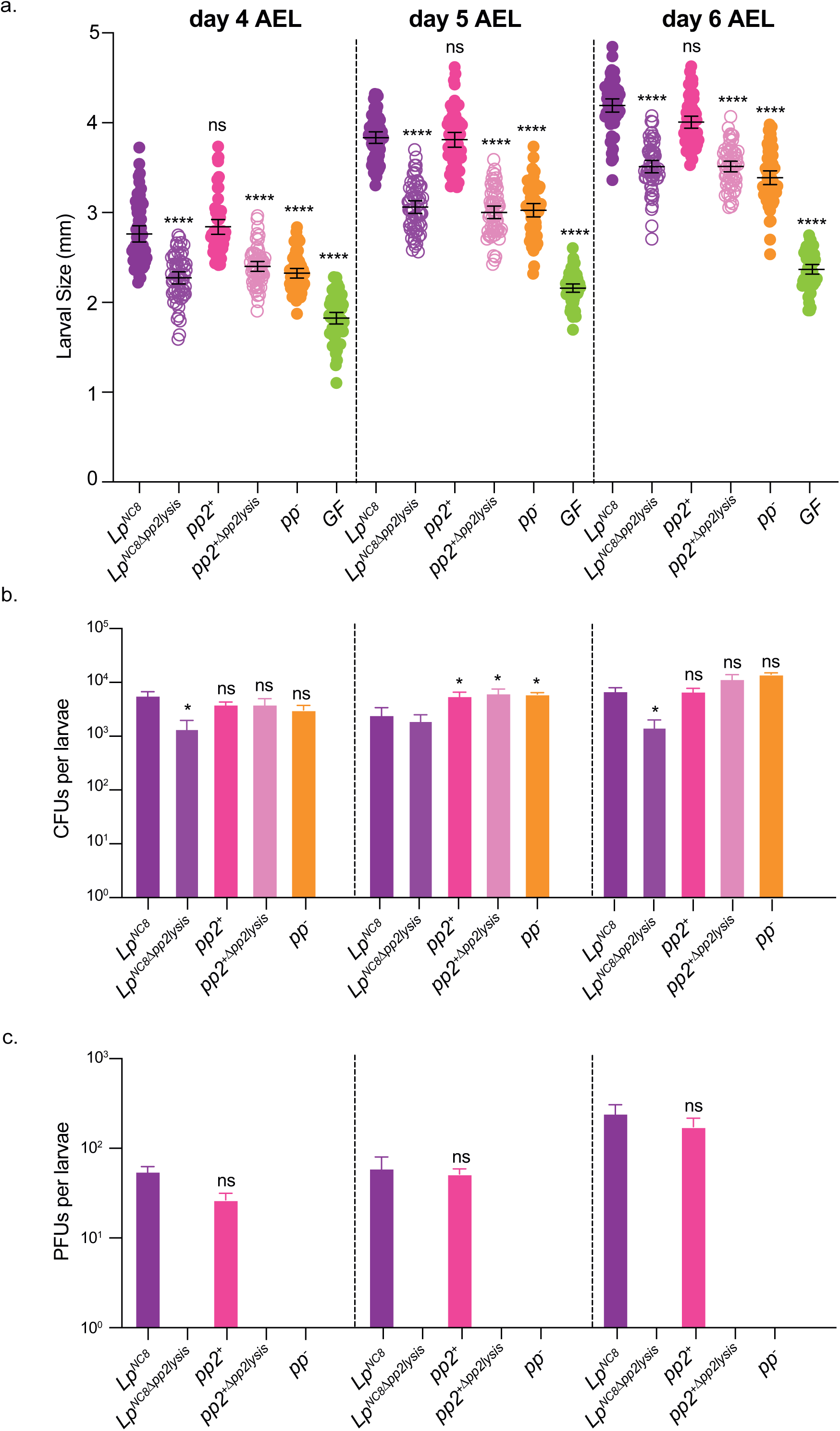
Prophage induced-lysis promotes drosophila growth. (a) Larval longitudinal length after inoculation with *Lp^NC8^*, *Lp^NC8^_Δpp2lysis_*, *pp2^+^*, *pp2^+^_Δpp2lysis_*, *pp^-^* strains and PBS for the germ-free condition. Larvae (n=60) were collected 4, 5 and 6 days after association and measured as described in the ‘Materials and methods’ section. Asterisks illustrate statistically significant differences with larval size of *Lp^NC8^* (p<0.05). Center values in the graph represent means and error bars represent 95% CI. Representative graph from one out of three independent experiments. Significance was determined using Kruskal-Wallis and Dunn’s multiple comparison tests. (b) Quantity of CFU associated with larvae at days 4, 5 and 6 after inoculation. Asterisks illustrate statistically significant difference with CFUs of *Lp^NC8^* associated larvae. Center values in the graph represent means and error bars represent SD. Representative graph from one out of three independent experiments. Significance was determined using Kruskal-Wallis and Dunn’s multiple comparison tests. (c) Quantity of PFU produced by bacteria associated with larvae at days 4, 5 and 6 after inoculation. Asterisks illustrate statistically significant difference with PFUs of *Lp^NC8^* associated larvae. Center values in the graph represent means and error bars represent SD. Representative graph from one out of three independent experiments. Significance was determined using Kruskal-Wallis and Dunn’s multiple comparison tests.

Strains capable of inducing prophage-mediated lysis - *Lp^NC8^*and *pp2⁺* - promoted thus significantly greater larval growth than lysis-deficient strains or prophage-free controls. These findings indicate that the enhanced growth-promoting effect of *Lp^NC8^* relies, at least partly, on pp2-mediated lysis and the concomitant release of symbiotic cues. These results establish that phage-induced lysis emerges as a central mechanism by which *Lp^NC8^* enhances host growth under nutritional stress.

### *Drosophila* intestinal physiology triggers prophage induction

Having established that bacterial lysis is required for *Lp*-mediated juvenile growth promotion in *Drosophila*, we next sought to identify what physiological parameter of the host intestine would trigger prophage induction, bacterial lysis, and consequently the release of symbiotic cues within extracellular vesicles. The *Drosophila melanogaster* gastrointestinal tract is organized into three main regions: the foregut, midgut, and hindgut^30^. Among these, the midgut is the principal site of host-microbe interactions, digestion and nutrient absorption playing a central role in intestinal homeostasis. Importantly, the midgut contains a specialized acidic compartment known as the copper cell region, which functionally resembles the mammalian stomach and maintains a regionalized low luminal pH^31^. In addition to acidity, the *Drosophila* gut deploys several mechanisms controlling microbial colonization while maintaining beneficial interactions with commensal bacteria. Accordingly to the luminal microbial load, intestinal epithelial cells activate conserved innate immune pathways, including the Imd and JAK/STAT pathways, leading to the production of antimicrobial peptides (AMPs) that restrict bacterial growth and contribute to gut homeostasis^30,32^. Furthermore, the *Drosophila* genome encodes several gut-expressed lysozymes that participate in the degradation of bacterial peptidoglycan and the regulation of microbial composition^33^. Together, the acidic environment of the midgut, the production of AMPs, and lysozyme activity create a physiological environment exerting strong selective pressures on gut-associated bacteria. We therefore hypothesized that these host-derived stresses contribute to bacterial cell-wall remodeling, prophage induction, lysis, and extracellular vesicle release in commensal bacteria such as *Lp*.

Taking advantage of the powerful *Drosophila* genetic toolkit, we tested the growth-promoting ability of *Lp^NC8^* in flies impaired in selected aspects of intestinal physiology in relation with host-microbe interactions. Specifically, we used flies deficient in gut lysozyme production (LysB-P^KO^)^34^, flies lacking major antimicrobial peptides (AMP-B^KO^)^35^, and flies in which differentiation of the acidic copper cell region was disrupted through enterocyte specific interference with the *labial* gene expression using a RNAi transgene under the control of the UAS/GAL4 system (*mex*>*lab*-IR)^36^ (*UAS-lab-IR*, BDSC#26753). We inoculated germ-free eggs from each fly line with *Lp^NC8^* and measured larval length at 4, 5, and 6 days after egg laying. As shown in Figure 7a, *Lp^NC8^*remained beneficial in individuals deficient for lysozyme production or antimicrobial peptides, as larvae associated with *Lp^NC8^* were significantly longer than their germ-free counterparts. In contrast, the growth-promoting effect of *Lp^NC8^* was strongly impaired in flies lacking the acidic copper cell region. At all-time points analyzed, larvae associated with *Lp^NC8^* in this genetic background displayed reduced growth compared to the corresponding genetic control (*mex>mCherry*), in which *Lp^NC8^* retained its full growth-promoting capacity (Fig. 7a).

**Fig. 7.**
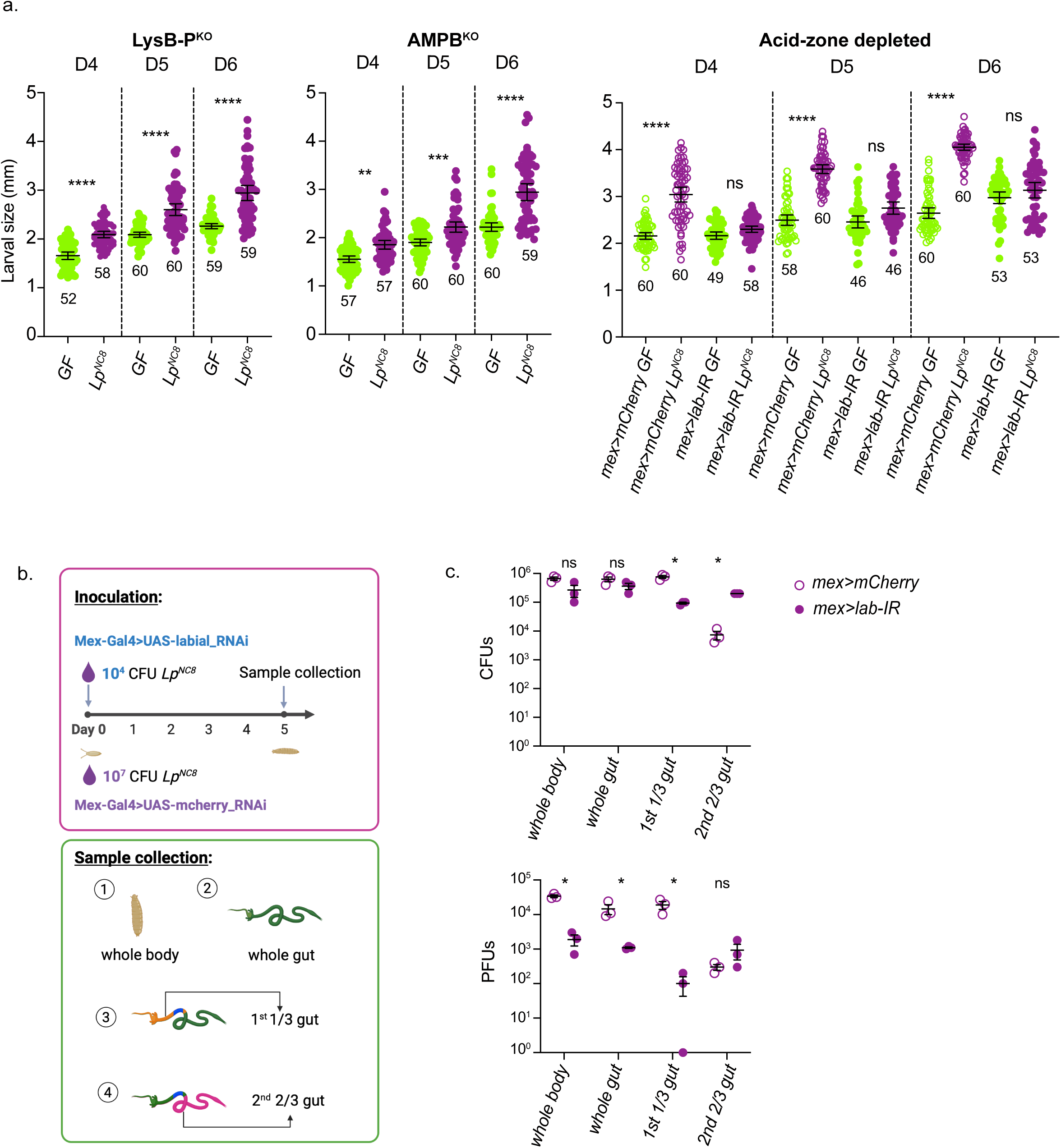
The Drosophila acidic intestinal region promotes prophage induction and phage release in *Lp^NC8^*. (a) Larval longitudinal length measurements of germ-free and *Lp^NC8^* associated larvae in different *Drosophila* genetic backgrounds impaired in host-microbe interaction mechanisms. Germ-free eggs from control flies, lysozyme-deficient flies (LysB-P^KO^), antimicrobial peptide-deficient flies (AMP-B^KO^), and flies lacking the acidic copper cell region through interference in *labi*al expression (*mex>lab-IR*) were inoculated with 10^7^ CFUs of *Lp^NC8^*. Larval longitudinal length was measured at 4, 5, and 6 days after egg laying (AEL) on low-protein fly food (7g/L of yeast). Asterisks represent statistically significant difference compared to *Lp^NC8^* (WT) larval size. Significance was determined using Kruskal-Wallis. ****P<0.0001. The bars in the graph represent mean and 95% CI. A representative graph from one out of three independent experiments is shown. For each condition n is indicated in the graph. (b) Experimental design used to quantify bacterial loads and phage production in larvae with or without the acidic midgut region (highlighted in blue). To obtain comparable bacterial colonization levels, *mex>mCherry* controls were inoculated with 10^7^ CFUs of *Lp^NC8^*, whereas *mex>lab-IR* individuals were inoculated with 10^4^ CFUs. Larvae were harvested 5 days after inoculation, and CFUs and PFUs were quantified from whole larvae, whole dissected guts, the anterior/middle midgut compartment containing the acidic region, and the posterior midgut compartment. (c) Quantification of CFUs and PFUs recovered from the indicated compartments. Adjusting the initial inoculum allowed comparable CFU levels between *mex>mCherry* and *mex>lab-IR* flies in whole larvae and dissected guts. The bars in the graph represent mean and 95% CI. A two-sided Mann-Whitney test was applied to perform pairwise statistical analysis between conditions. Asterisks illustrate statistically significant differences (p<0.05).

Based on these results, we hypothesize that passage through the gut’s acidic region triggers prophage induction, leading to phage release, which we use here as a proxy for *in vivo* bacterial lysis and extracellular vesicle production. Previous studies have shown that the acidic zone exerts bactericidal activity on gut-associated microbes^37^, with commensal bacteria accumulating in the anterior midgut in control individuals while higher total intestinal bacterial loads with a striking increase in posterior midgut bacterial loads were detected in *labial* mutants lacking this compartment compared to their genetic controls. To specifically assess whether the acidic zone influences lysis and phage release independently of bacterial abundance, we established experimental conditions yielding comparable CFU levels in both genetic backgrounds. To do so, we inoculated *mex>mCherry* controls and *mex*>*lab-RNAi* individuals with different initial concentrations of *Lp^NC8^* and quantified midgut bacterial loads 5 days after egg laying (Fig. 7b). As shown in Supplementary Figure 5, inoculation of *mex>mCherry* controls with 10^7^ CFUs and *mex*>*lab-RNAi* individuals with 10^4^ CFUs resulted in comparable bacterial loads in the midgut 5 days after inoculation. Next, we determine the number of PFUs produced by comparable bacterial loads in larvae with or without the acidic gut region. Larvae were inoculated with 10^7^ CFUs of *Lp^NC8^* for *mex>mCherry* controls and 10^4^ CFUs for *mex*>*lab-RNAi* individuals (Fig. 7b). Larva were harvested 5 days after inoculation, and both CFUs and PFUs were quantified from whole larvae, whole dissected midguts, the anterior/middle part of the midgut containing the acidic region, and the posterior midgut compartment. Figure 7c shows that, by adjusting the initial inoculum, we obtained comparable CFU levels in whole larvae and whole dissected midguts between *mex>mCherry* controls and *mex>lab-RNAi* individuals. In *mex>lab-RNAi* individuals, CFU counts in the anterior midgut compartment, which includes the acidic region, were slightly decreased while strongly increased in the posterior midgut. In contrast, *in vivo* PFU quantification revealed a consistent and marked reduction in phage production in flies lacking the acidic gut region, a decrease observed across all compartments analyzed. Together, these results identify the acidic region of the *Drosophila* midgut as a key intestinal physiological parameter promoting *in vivo* prophage induction, bacterial lysis in *Lp^NC8^* and support to host growth despite nutritional stress.

## DISCUSSION

Understanding how commensal bacteria generate, package, and deliver bioactive molecules to their hosts remains a central challenge in microbiome biology. Here, we identify prophage-induced lysis as a previously unappreciated physiological strategy that enables a symbiont that supports adaptive host growth to release host-active cues. Using *Lactiplantibacillus plantarum* NC8 (*Lp^NC8^*), a commensal that promotes *Drosophila* juvenile growth under nutritional stress, we show that its resident prophage, pp2, is not an inert genomic element but an environmentally responsive module essential for host adaptation to nutritional stress. pp2 induction activates a holin-lysin lytic program that produces phage virions and extracellular particles, and this controlled lysis is required for *Lp*-mediated growth promotion *in vivo*. These findings establish a mechanistic link between bacterial lysis and microbe-driven adaptation of animal juvenile growth to nutritional stress.

Systematic characterization of *Lp*^NC8^ prophages revealed pp2 as a stress-responsive element capable of executing full lytic cycles. Exposure to DNA-damaging agents triggered robust pp2 activation, leading to holin-lysin-dependent membrane and cell wall permeabilization, release of infectious virions, and the generation of abundant extracellular vesicles with their morphology and origin consistent with membrane-derived structures reminiscent of vesicles produced by diverse Gram-positive and Gram-negative bacteria^27,38,39^. Importantly, these vesicles contain LTAs, a major Gram-positive cell envelope component and a symbiotic cue previously identified as essential for *Lp*-mediated juvenile growth promotion^9,10^, thereby providing a mechanistic framework linking vesicle release to delivery of host-active bacterial molecules. Prior work in *Bacillus subtilis* and *Staphylococcus aureus* has similarly implicated phage-mediated cell envelope rupture in vesicle biogenesis^15,26,27^. Our data support a model in which pp2-driven lysis contributes to the formation of such particles, which may facilitate the trafficking, stability, or epithelial targeting of bacterial molecules within the gut^40–42^. A major next step will be to characterize their molecular cargo, and to map their production dynamics *in vitro* and *in vivo*.

The discovery that pp2-mediated lysis is necessary and sufficient for the growth-promoting activity of *Lp*^NC8^ represents a conceptual shift in how beneficial bacteria deliver symbiotic cues. Several *Lp*-derived molecules, including peptidoglycan^8^, D-alanylated lipoteichoic acids^9,10^, and r/tRNAs^11^, support juvenile growth, yet their route of delivery to host tissues has remained obscure. Most of these cues are intracellular or embedded in the bacterial cell envelope, and the architecture of the midgut, including the peritrophic matrix, poses substantial physical barriers to their release and epithelial access^19^. Our findings suggest that prophage induction provides a major pathway by which structurally diverse cues escape the bacterial cell and enter the intestinal milieu, enabling them to reach host targets despite anatomical constraints. Thus, lysis emerges not as a detrimental event but as a physiologically integrated mechanism of interkingdom signalling.

The responsiveness of pp2 to DNA damage is consistent with canonical mechanisms of SOS-dependent prophage induction^23^. Importantly, our *in vivo* data now indicate that host intestinal physiology itself contributes to triggering prophage induction. We show that disruption of the acidic copper cell region of the *Drosophila* midgut strongly impairs *Lp^NC8^*-mediated growth promotion and is accompanied by a marked reduction in phage release despite high bacterial loads. These findings identify the acidic midgut compartment as a key host physiology parameter promoting prophage activation *in vivo*. The *Drosophila* gut combines multiple bacterial stressors, including low pH, antimicrobial peptides, and lysozyme activity, all of which may contribute to bacterial envelope stress and DNA damage responses^43^. Previous work demonstrated that the acidic region exerts potent bactericidal activity toward gut-associated microbes^37^, and our data extend this concept by showing that this physiological stress also modulates prophage activity and bacterial lysis dynamics *in vivo*. Related phenomena have been observed in mammalian systems, where prophage induction shapes microbial community composition and modulates host responses such as bile acid metabolism^18,44,45^. Our work expands this framework by demonstrating that host-derived physiological constraints can directly regulate prophage-mediated release of beneficial bacterial cues for host adaptation to nutritional stress. *In vivo*, the timing and magnitude of pp2 activation correlate strongly with larval growth promotion, supporting a model in which environmental stress in the intestine modulates prophage activity, which in turn regulates the release of *Lp*-derived symbiotic cues. This raises the intriguing possibility that the host actively tunes bacterial secretory and lytic dynamics through intestinal physiology. In this context, the acidic region of the gut may function not only as an antimicrobial barrier but also as a spatially restricted regulator of beneficial host-microbe communication.

From the bacterial perspective, maintaining an inducible prophage may appear costly, yet pp2 likely confers ecological advantages at the population level. Lysis of a subpopulation can release intracellular metabolites, membrane fragments, and phage virions that aid surviving clonal members, an effect that may be particularly advantageous in nutrient-poor environments such as the undernourished gut. Importantly, pp2-driven lysis also liberates symbiotic cues that promote host juvenile growth, thereby improving the nutritional landscape in which *Lp* thrives. Consistent with this mutually reinforcing dynamic, we previously showed that *Drosophila* larvae excrete “maintenance factors” that fortify the food substrate and sustain their commensal partner over time^37^, suggesting that host benefits feed-back to stabilize bacterial populations. Although prophages are frequently viewed through the lens of virulence, lysogenic conversion, or microbial competition^46–48^, pp2 instead operates as an inducible symbiotic module that links environmental stress to the controlled release of host-active factors. Our findings therefore support a broader model in which host physiology, bacterial stress responses, prophage induction, and extracellular vesicle release are mechanistically interconnected.

In conclusion, we reveal prophage induction as a mechanism by which a commensal bacterium promotes host growth. By coupling environmental stress responses to phage activation and release of symbiotic molecules, *Lp*^NC8^ leverages an inducible prophage to support host development under nutritional challenge. These findings expand the conceptual framework of beneficial host-microbe interactions, highlighting prophages as dynamic and integral participants in symbiosis and identifying host intestinal physiology as a direct regulator of bacterial lysis dynamics and vesicle-mediated symbiotic communication.

## METHODS

### Drosophila diets, stocks and breeding

Drosophila stocks were cultured as described previously^8^. Briefly, flies were kept at 25°C with 12/12-hour dark/light cycles on a yeast/cornmeal medium containing 50 g.l^−1^ of inactivated yeast. The poor-yeast diet was obtained by reducing the amount of inactivated yeast to 7 g.l^−1^. Germ-free stocks were established as described previously^49^. Axenicity was routinely tested by plating serial dilutions of animal lysates on nutrient agar plates. *Drosophila y,w* flies were used as the reference strain in this work The following *Drosophila melanogaster* lines were used in this study: *y,w*, *LysB-P^KO^*^34^, *AMP-B^KO^*(*w; AttCMi, DroAttSK2, DptSK1; AttDSK1*)^35^, *mex-Gal4*^36^, *UAS-labial-IR*, and *UAS-mCherry-IR*. *LysB-P^KO^* and *AMP-B^KO^* lines were obtained from the laboratory of Bruno Lemaitre. *mex-Gal4* line was obtained from the laboratory of Irene Miguel-Aliaga. *UAS-labial-IR* (#26756), and *UAS-mCherry-IR* (#35785) lines were obtained from Bloomington stock center. *mex>lab-IR* progeny were generated by crossing *mex-Gal4* driver flies with *UAS-labial-IR* flies, while *mex>mCherry-IR* progeny were generated by crossing *mex-Gal4* driver flies with *UAS-mCherry-IR* flies and used as genetic controls.

### Bacterial strains and growth conditions

The strains used in this study are listed on Extended Data Table 1. *Escherichia coli* strains were grown at 37°C in LB medium with agitation. *L. plantarum* strains were grown in static conditions in MRS medium at 37°C, unless stated otherwise. Erythromycin antibiotic was used at 5 μg.ml^−1^ for *L. plantarum* and 150 μg.ml^−1^ for *E. coli*. Chloramphenicol antibiotic was used at 25 μg.ml^−1^.

### Construction of prophage deleted strains in *Lp^NC8^* using pG+host9

Markerless deletions of individual or combinatorial prophages were generated by homology-directed recombination using a double-crossing over strategy, as previously described^9^. Briefly, the 5′ and 3′ terminal regions of each prophage were PCR-amplified from *Lp^NC8^*genomic DNA using Q5 High-Fidelity 2X Master Mix (New England Biolabs). Primers were designed to contain overlaps with the temperature-sensitive vector pG+host9^50^ to enable Gibson assembly of the deletion constructs. The following primer pairs were used to amplify homology arms: OL13/OL14 and OL15/OL16 for pp1, OL19/OL20 and OL21/OL22 for pp2, and OL25/OL26 and OL27/OL28 for pp3 (see Extended Data Table 2). Recombinant plasmids were transformed into *Lp^NC8^*, and integration and excision events were selected through temperature shifts as described previously. Successful deletions were confirmed by colony PCR using primer pairs OL17/OL18 (pp1), OL23/OL24 (pp2), and OL29/OL30 (pp3).

### Construction of *recA* and pp2 lysis module deletions in *Lp^NC8^*using pG+host9

Markerless deletions of *recA* and of the pp2 lysis module (*nc8_0567–nc8_0568*) were generated by homology-directed recombination using a double-crossing over strategy, as previously described (Matos et al., 2017). Briefly, the 5′ and 3′ terminal regions flanking each targeted locus were PCR-amplified from *Lp^NC8^*genomic DNA using Q5 High-Fidelity 2X Master Mix (New England Biolabs). Primers were designed with overlapping sequences compatible with the temperature-sensitive plasmid pG+host9^50^ to enable Gibson assembly of the deletion constructs. Homology arms were amplified using primer pairs OL01/OL02 and OL03/OL04 for *recA*, and OL07/OL08 and OL09/OL10 for *nc8_0567–nc8_0568* (Table S2). Following transformation into *Lp^NC8^*, integration and excision events were selected through temperature shifting. Deletions were verified by PCR using primer pairs OL05/OL06 (*recA*) and OL11/OL12 (*nc8_0567–nc8_0568*).

### Conditions of prophage induction and plaque-forming unit quantification

Overnight cultures of the tested *Lp* strains (*Lp^NC8^*, *pp1^+^*, *pp2^+^*, *pp3^+^*, and derivatives) were back-diluted 1:100 into fresh MRS medium and incubated at 28 °C until early exponential phase. Prophage induction was triggered by supplementing cultures with mitomycin C (MMC; 2 µg.mL^-1^), ciprofloxacin (0.4 mg.mL^-1^), or polymyxin B (15 mg.mL^-1^). Samples were collected at the indicated time points following treatment. For colibactin-mediated induction, overnight cultures of *E. coli pks^+^*and *pks^-^*strains grown in LB medium supplemented with chloramphenicol (25 µg.mL^-1^) at 37 °C were washed with sterile 1x PBS to remove residual medium, then back-diluted 1:100 into BHI medium together with the *Lp* strain. Co-cultures were incubated at 28°C and sampled at the indicated times. For plaque assays, culture or co-culture samples were filtered through 0.22 µm MES-membrane filters, and filtrates were serially diluted in SM (10 mM Tris-HCl pH 7, 10 mM NaCl, 10 mM MgSO_4_) buffer. Dilutions were spotted onto lawns of the appropriate indicator strain embedded in MRS top agarose (0.5%) supplemented with CaCl_2_ (10 mM) and MgCl_2_ (10 mM). Plates were incubated at room temperature for 24 h, after which plaque-forming units (PFUs) were enumerated.

### Optical microscopy

#### Snap shots imaging for bacterial viability

Bacterial cell viability was assessed using the LIVE/DEAD® BacLight™ Bacterial Viability Kit (Thermo Fisher Scientific). Cultures from both mitomycin C (MMC)-treated and untreated groups were harvested by centrifugation at 6,000 × *g* for 10 min. Supernatants were discarded, cell pellets were gently washed twice with 0.85% NaCl, and resuspended in 0.85% NaCl. SYTO 9 and propidium iodide (PI) stains were mixed in a 1:1 (v/v) ratio according to the manufacturer’s instructions. The dye mixture was added to the bacterial suspension at a ratio of 3 μL of dye mix per 1 mL of cell suspension. The stained suspensions were incubated in the dark at room temperature for 20 min, and placed on slides for observation. Images were acquired using Leica DM 6000 equipped with appropriate lasers and filters. The acquired images were analyzed using Fiji software^51^. The numbers of green-fluorescent cells (live) and red-fluorescent cells (dead) were counted manually. The percentage of viable cells was calculated as follows: Viability (%) = [Number of green cells / (Number of green cells + Number of red cells)] × 100%.

#### Snap shots imaging for morphology analyses

Exponentially growing bacterial cells were washed with 1 × PBS (6,000 x *g*, 5 min) and resuspended in PBS. Cells were incubated with FM4-64 (10 μg/mL) (Invitrogen by Thermo Fisher Scientific) in the dark for 5 min at room temperature. Bacterial cells were then placed on top of PBS-agarose (1%) pads on microscopy slides and covered by coverslips. Slides were visualized using a Nikon TiE microscope, equipped with the perfect focus system (PFS, Nikon), and fitted with an Orca-CMOS Flash4 V2 camera with a 100 × 1.45 objective. Images were collected using NIS-Elements (Nikon) and processed with Fiji software^51^. Cell length quantification was performed with the MicrobeJ plugin^52^ in combination with masks generated with OmniPose^53^. For visualization purpose, brightness and contrast were adjusted for phase contrast and FM4-64 signals.

#### Time-lapse imaging

Overnight precultures of bacteria were back diluted 1:100 into fresh MRS and incubated at 28 °C for 6h. Bacterial cells were placed on top of MRS-agarose pad (1%) supplemented with FM4-64 (10 μg.mL^-1^), with or without MMC (2 µg.mL^-1^). Slides observation and images acquisition were performed at 30°C through a thermostated chamber, with the same Nikon microscope setup as above. Phase contrast images were captured every 30 min and FM4-64 images every hour. Time lapse images were processed with Fiji software^51^, and x/y drift over time was corrected using the HyperStackReg plugin. For visualization purpose, brightness and contrast were adjusted for phase contrast and FM4-64.

##### Lysozyme assay

Cultures of *Lp* strains were induced with MMC as described above. After 24 h, cells were pelleted to initiate the lysozyme digestion assay, while the culture supernatants were filtered (0.22 µm) to quantify PFUs released during prophage-induced lysis. Pelleted cells were resuspended in fresh MRS medium and incubated for 1 h with lysozyme (5 mg.mL^-1^). Following incubation, samples were centrifuged and the resulting supernatants were collected. PFUs in these supernatants were quantified by plaque assay to determine the number of phage particles liberated by lysozyme-mediated cell wall digestion, representing phages that remained intracellular after MMC induction and were not released during prophage-induced lysis.

##### Transmission electron microscopy

Overnight bacterial cultures were back-diluted 1:100 into fresh MRS medium and incubated at 28°C for 6 h. Cultures were then either induced with mitomycin C (MMC; 2 µg.mL^-1^) and incubated for an additional 24 h or maintained without MMC for 24 h as controls. Cells were harvested by centrifugation (6000 g, 5 min), washed once with sterile 1x PBS, and resuspended in PBS. For grid preparation, 5 µL of the suspension was applied to 200-mesh copper grids coated with carbon-stabilized formvar films (Electron Microscopy Sciences) and allowed to adsorb for 1 min. Excess liquid was removed using filter paper. Grids were negatively stained by floating them on a drop of 1% (w/v) aqueous uranyl acetate for 30 s, followed by blotting and air-drying. Samples were imaged using a JEOL 1400 Flash transmission electron microscope.

### Extracellular vesicles

#### Isolation and purification

Overnight cultures of the tested *Lp* strains were back-diluted 1:100 into fresh 0.5 L MRS medium and incubated at 28°C until early exponential phase. Prophage induction was triggered by supplementing cultures with MMC. The number of bacteria was determined using CFU counting. Bacteria was spun down (6000 x g, 10 min) and the supernatant was filtered through 0.22 µm filters. Filtrates were ultracentrifuged at 150 000 x g, for 3 h, at 4°C using Sorvall WX+ centrifuge and T-647.5 Fixed Angle Rotor (ThermoFisher Scientific). Pellets were re-suspended in 25 mM HEPES with 0.9 NaCl. Crude EVs were purified using Size Exclusion Chromatography (SEC) qEV columns (IZON, New Zealand), as described previously^54^. Six 400 µl fractions were collected. Size and concentration of particles were analyzed with Zetasizer Ultra (Malvern Panalytical, UK). Zetasizer working ranges were of: concentration measurement range of 1×10^8^ to 1×10^12^ particles/mL and diameter of 0.3nm - 15µm. Each measurement was done using General Mode and in triplicate. Measurement quality and data analysis were performed using Malvern ZS Xplorer software (version 3.3.1.5).

#### Transmission electron microscopy

TEM microcopy was performed as previously published^55^. 5 µl of EV fraction 1.2 of each sample was deposited on the formvar carbon coated copper grid, mesh 400 and incubated for 2 min. The excess fluid was drained with filter paper. 4 µl of 1% uranyl acetate was added, left for 1 min and drained. Images were acquired with Jeol JEM-1200EX with accelerating voltage 80 KV and Veleta camera.

#### Western blot analysis

5 µl of x4 concentrated Laemmli buffer was added to 15 µl of each fraction and boiled for 5 min at 96°C. Samples were applied to 12% gels (BioRad, USA) which were then used for Sodium Dodecyl Sulfate Polyacrylamide Gel Electrophoresis for 1.5 h at 120 V. Gel was blotted to the PVDF membrane (0.45 µm pore size, Sigma-Aldrich), blocked with SuperBlock buffer for 120 min at room temperature (ThermoFisher Scientific) and incubated (2 h, at room temperature) with 1:100 primary anti-LTA (MA1-7402, Invitrogen) antibodies in Tris-HCl buffer, pH 7.0, 0.1% Tween 20 (TBS-T). Next, after washing, they were incubated (1 h, at room temperature) with 1:30 000 secondary anti-mouse IgG (A3562, Sigma-Aldrich) antibodies conjugated with alkaline phosphatase in TBS-T. Blot was developed with NBT/BCIP (Roth, Germany). The reaction was stopped with water wash. Blot was documented with ChaemiDoc MP Imaging System (BioRad). 25 µg of LTA from Staphylococcus aureus (L2515, Sigma-Aldrich) was used per well as positive control. 5 µl per well of Pierce™ Pre-stained Protein MW Marker (ThermoFisher Scientific) was used.

#### Larval size measurements and quantification of bacterial and phage loads

Larval growth was assessed using a gnotobiotic *Drosophila melanogaster* assay as previously described^8^. Axenic adults were placed in sterile breeding cages overnight to lay eggs on poor-yeast diet. Freshly collected axenic embryos were transferred the following morning and seeded in pools of 40 onto 55-mm petri dishes containing sterile poor-yeast diet. Bacterial suspensions (1×10^7^ CFU, unless otherwise specified) or sterile phosphate-buffered saline (PBS) were applied homogeneously onto both the food substrate and embryos. Plates were incubated at 25°C, and larvae were collected at day 6 post-inoculation unless indicated otherwise. Individual larvae were rinsed, mounted, imaged under a stereomicroscope, and larval longitudinal length was quantified using ImageJ^51^. For quantification of bacterial and phage loads, 40 larvae were collected into sterile 100-µm cell strainers and surface-sterilized with 100% ethanol, followed by two rinses in sterile 1x PBS. Larvae were transferred to sterile microcentrifuge tubes and homogenized in 1x PBS with glass beads at 4600 rpm for 30 s (three cycles) using a Precellys 24 tissue homogenizer (Bertin Technologies). Homogenates were serially diluted and plated on MRS agar, and plates were incubated at 28°C overnight to determine colony-forming units (CFUs). To quantify phage particles, supernatants were serially diluted and spotted onto lawns of the appropriate indicator strain embedded in MRS top agarose (0.5%) supplemented with 10 mM CaCl_2_ and 10 mM MgCl_2_. Plates were incubated at room temperature for 24 h prior to plaque-forming unit (PFU) enumeration.

#### Quantification of bacterial and phage loads in dissected Drosophila midgut compartments

Axenic virgin females carrying *mex-GAL4* were crossed under sterile conditions with axenic males carrying *UAS-labial-IR*. Flies were allowed to mate and lay eggs overnight. Freshly collected axenic embryos were transferred the following morning in pools of 40 onto 55-mm Petri dishes containing sterile poor-yeast diet. To equalize bacterial loads between fly genotypes, embryos from the *mex>Lab-IR* cross were inoculated with 1 × 10^4^ CFU of *Lp^NC8^*. As a genetic control, axenic embryos obtained from crosses between *mex-GAL4* virgins and *UAS-mCherry-IR* males (*mex>mCherry-IR*) were inoculated with 1 × 10^7^ CFU of *Lp^NC8^*. Larvae were collected 5 days post-inoculation for subsequent analyses. For whole-body analysis, pools of 30 larvae were washed in 70% ethanol and rinsed twice in sterile PBS. For gut dissections, larvae were dissected in sterile 1x PBS. Whole gut samples included the digestive tract from the proventriculus to the hindgut, excluding Malpighian tubules. Dissected guts were further subdivided into two regions: (i) an anterior fraction corresponding to approximately the first third of the gut, including the foregut, anterior midgut, and acidic copper cell region; and (ii) a posterior fraction corresponding to the remaining two-thirds of the gut, including the posterior midgut and hindgut. Whole larvae, whole guts, or dissected gut regions were transferred into sterile microcentrifuge tubes and homogenized in 800 μL sterile PBS containing glass beads using a Precellys 24 tissue homogenizer (Bertin Technologies) at 4,600 rpm for 30 s over three cycles. CFU and PFU quantifications were then performed as described above.

#### Statistical analysis

Data representation and analysis was performed using Graphpad PRISM 10 software (www.graphpad.com). Three to five biological replicates were used for all experiments performed in this study in order to ensure representativity and statistical significance. All samples were included in the analysis. Experiments were performed without blinding. A two-sided Mann–Whitney test was applied to perform pairwise statistical analyses between conditions. A Kruskal-Wallis test was applied to perform statistical analyses between multiple (n>2) conditions followed by Dunn’s test that corrects for multiple comparisons.

## Declaration of generative AI and AI-assisted technologies in the writing process

During the preparation of this work, the authors used ChatGPT in order to polish the language of the manuscript. After using this tool, the authors reviewed and edited the content as needed and take full responsibility for the content of the publication.

## Acknowledgments

The authors thank Marie-Agnes Petit, Thomas Henry, Filipe De Vadder and Jacques Montagne for helpful discussions. Eric Oswald for sending colibactin producing strains. Chloé Exbrayat-Héritier from Centre Technologique des Microstructures EZUS LYON - Université Claude Bernard Lyon 1 for transmission electron microscopy experiments. PLATIM and Arthro-tools platforms from SFR Bioscience (UAR3444/US8) for providing imaging and Drosophila facilities. C.Z. was funded by Chinese Scholarship Council through the PROSFER program. The work was funded by seed money from ENS Lyon and FINOVI foundation and ANR grant (ANR-23-CE20-0004-01) awarded to R.C.M. The laboratory of F.L. was supported by “Fondation pour la Recherche Médicale” (Equipe FRM DEQ20180339196) for this work.

## Author contributions

Conceptualization: FL, RCM

Methodology: CZ, SM, CIR, AR, RCM

Formal analysis: CZ, SM, AR, FL, RCM

Investigation: CZ, SM, QP, SV, HA, NL, AR

Writing – original draft: FL, RCM

Writing – review and editing: CZ, SM, QP, SV, HA, NL, CIR, MZ, AR, CG, FL, RCM

Visualization: CZ, SM, AR, RCM

Supervision: FL, RCM

Funding acquisition: FL, RCM

## SUPPLEMENTARY DATA

**Supplementary Fig. 1.**
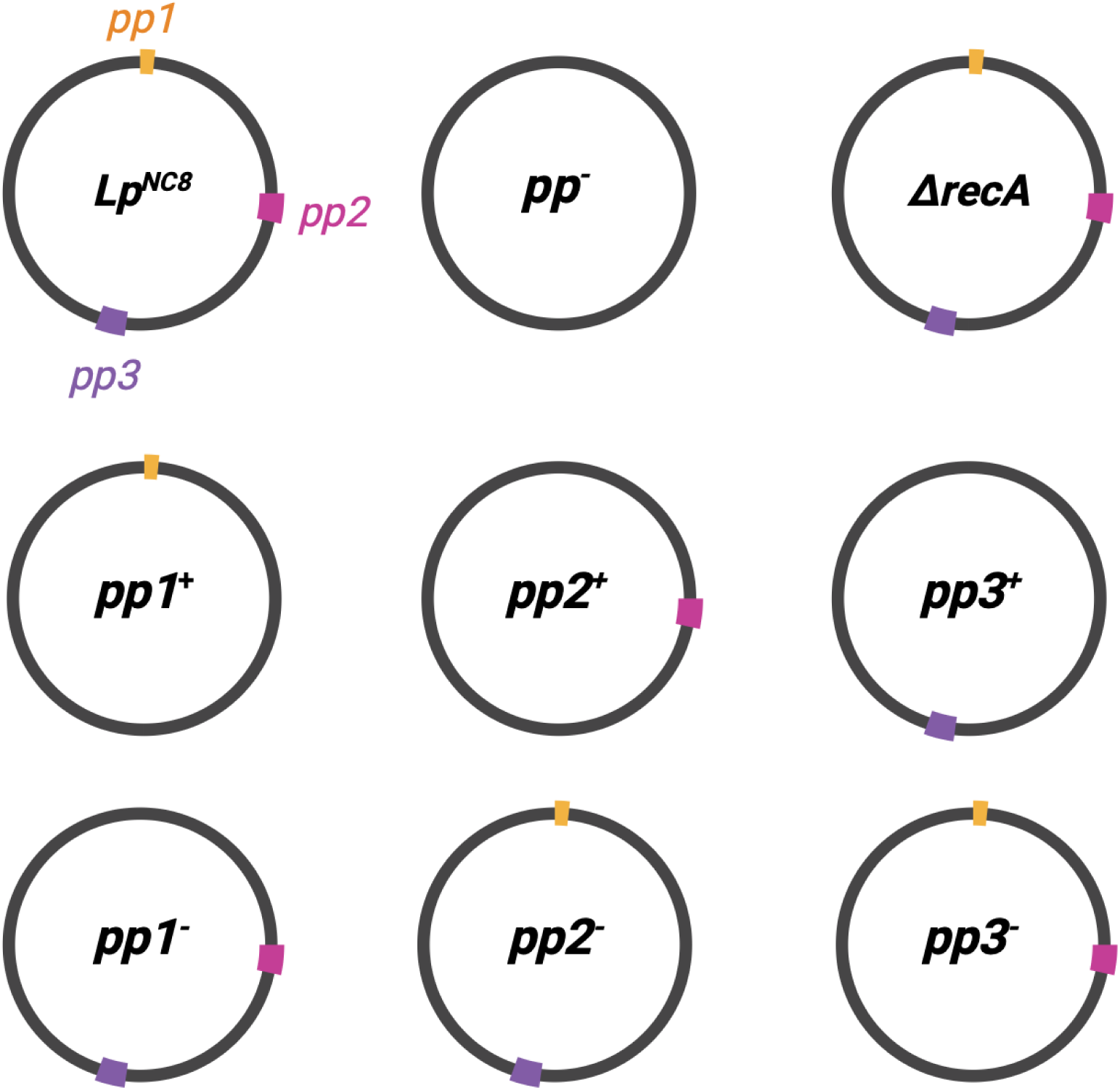
Isogenic *Lp^NC8^* strains generated in this study. The wild-type *Lp^NC8^* genome contains three integrated prophages (*pp1*, *pp2*, *pp3*). To dissect their individual contributions, we constructed monolysogenic strains in which only one prophage remains (*pp1^+^*, *pp2^+^*, *pp3^+^*), as well as the corresponding single-deletion strains (*pp1^-^*, *pp2^-^*, *pp3^-^*) used for superinfection assays. A prophage-free derivative was obtained by sequential removal of all three prophages (*pp^-^*). Finally, a Δ*recA* strain was also built to prevent SOS-dependent prophage induction.

**Supplementary Fig. 2.**
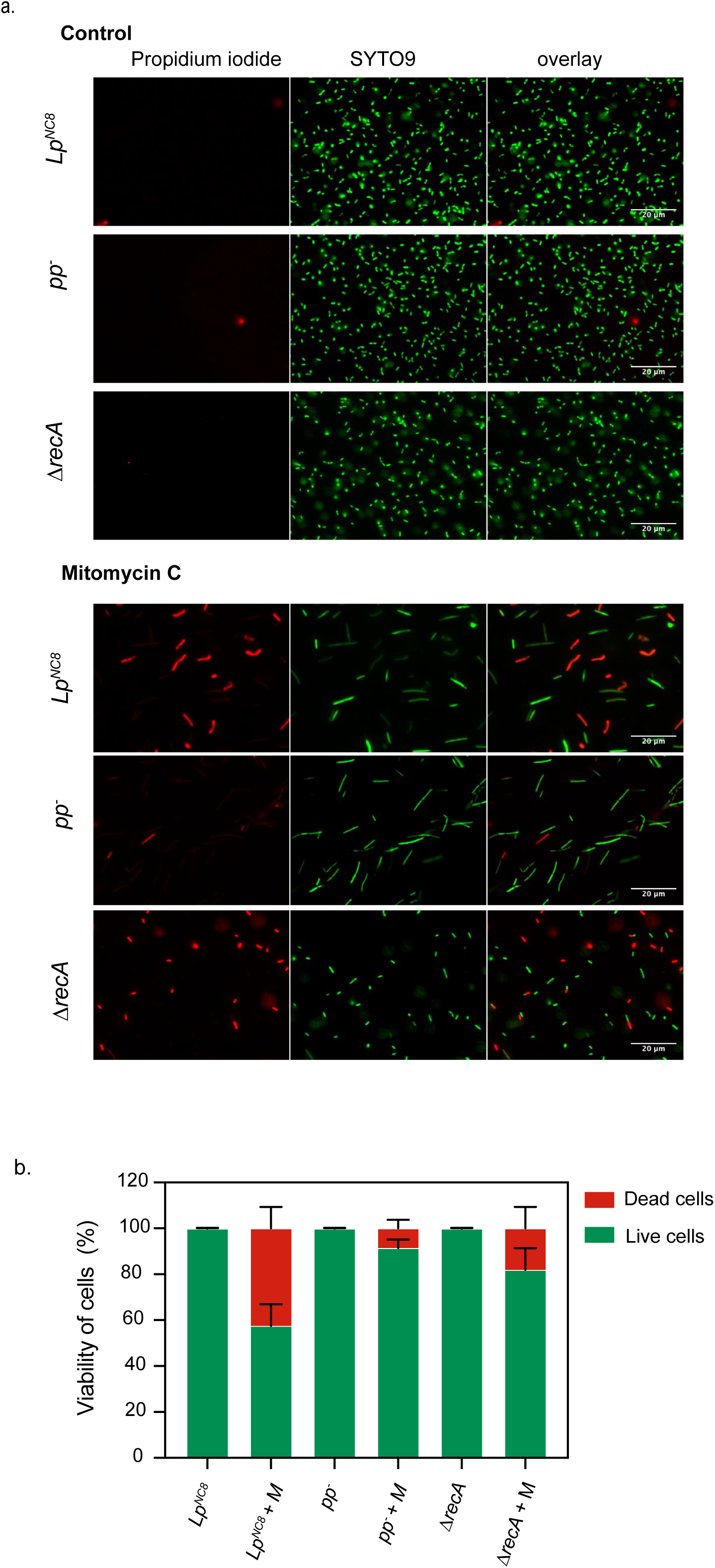
Cell viability. The LIVE/DEAD® BacLight™ Bacterial Viability Kit was used to quantify live/dead cells from *Lp^NC8^*, *pp-*, *ΔrecA* strains in presence or absence of mitomycin C. This kit distinguishes live from dead bacteria based on membrane integrity, using two nucleic-acid binding dies: SYTO9 (green) and propidium iodide (red). SYTO9 labels all cells while propidium iodide selectively penetrates and stains membrane-compromised cells; because propidium iodide displaces SYTO 9 upon DNA binding, viable cells fluoresce green whereas dead fluoresce red. (a) Representative images and (b) percentage of live or dead cells 24 hours after mitomycin C induction.

**Supplementary Fig. 3.**
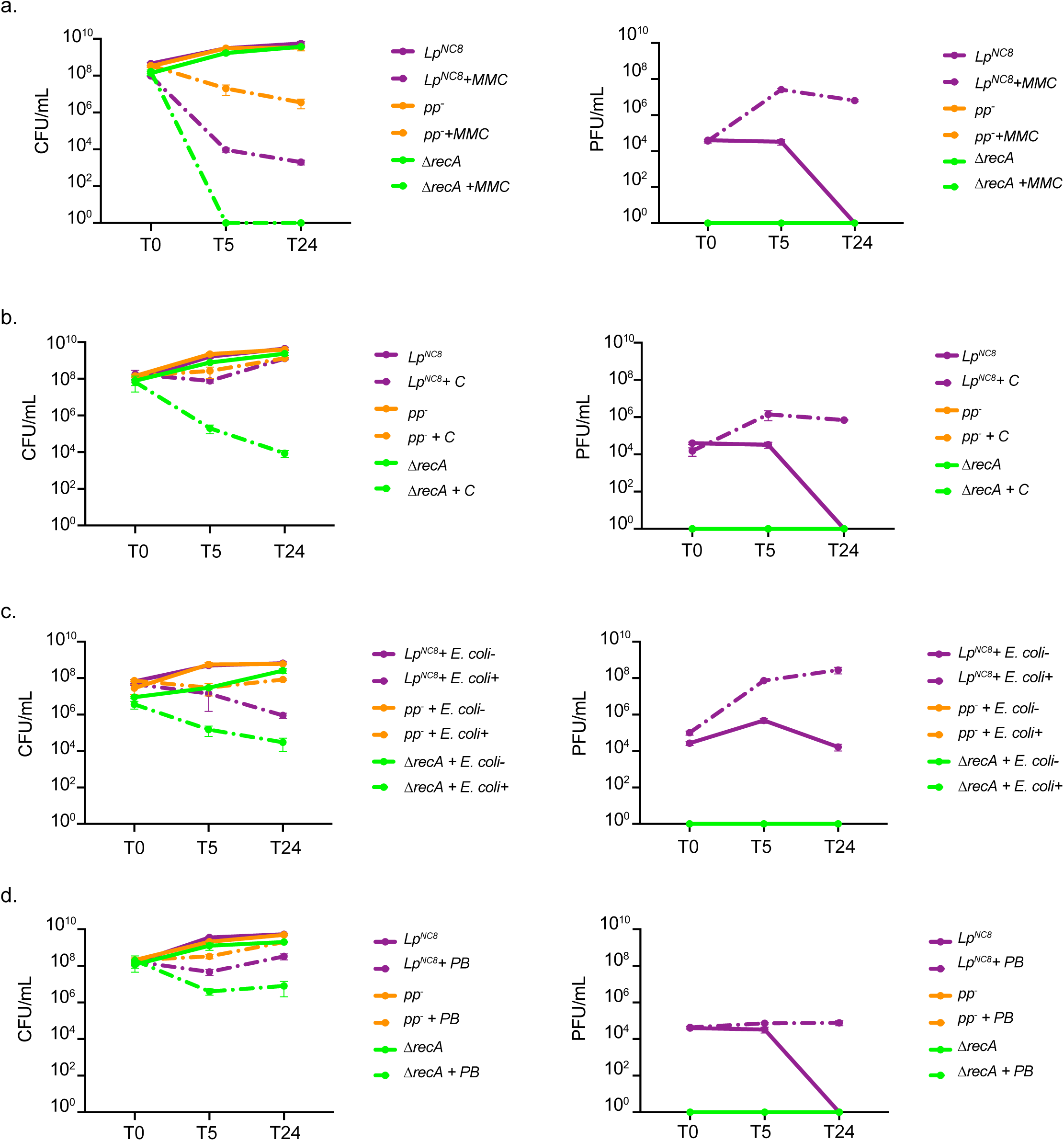
Prophage induction dynamics in presence of different stressors. CFUs *vs* PFU representation for *Lp^NC8^*, *pp*^-^ and *ΔrecA* strains upon different stressors supplementation (a) Mitomycin C (MMC), (b) ciprofloxacin (C), (c) colibactin (produced by *E. coli pks^+^)* and (d) polymyxin B (PB). Exponentially growing cultures were induced with mitomycin C. T0 corresponds at the time of mitomycin C supplementation. Cells and supernatants were harvested at different timepoints after induction and analyzed for CFUs and PFUs on a lawn of *pp^-^* strain, respectively.

**Supplementary Fig. 4.**
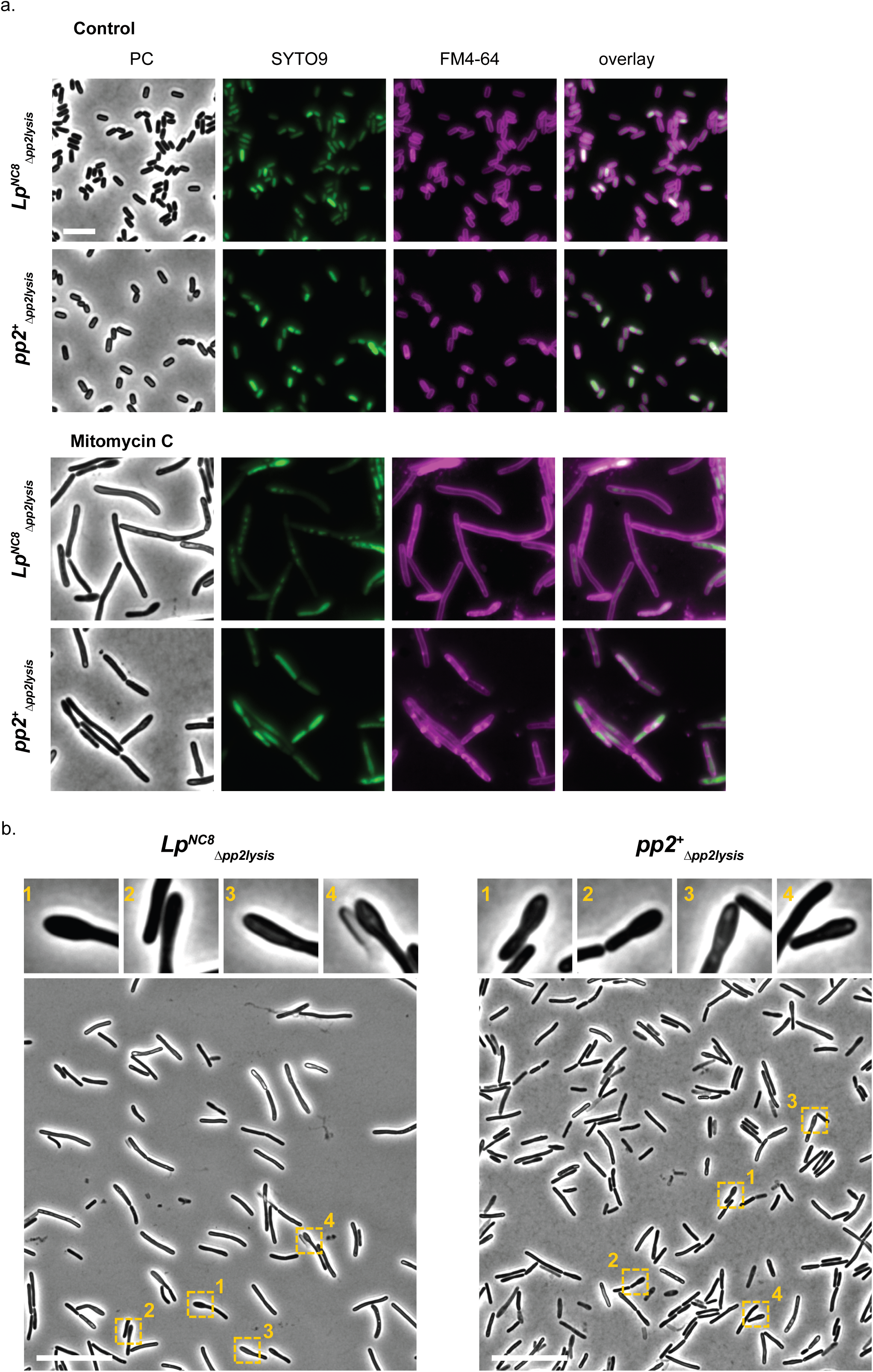
Loss of the pp2 lysis module blocks bacterial lysis and alters cell morphology. (a) Fluorescence microscopy of *Lp^NC8^*and *pp2^+^* lysis module deletions strains. Representative images of *Lp* cultures (*Lp^NC8^* and *pp2^+^*) treated or not with mitomycin C, and dyed with FM4-64 (false-colored in magenta) to stain membranes and SYTO9 (false-colored in green) to stain DNA. PC stands for phase contrast. Scale bars, 5 µm. (b) Phase contrast close-ups of spoon-shaped cells observed in *Lp^NC8^* and *pp2^+^* cultures upon induction with mitomycin C. Scale bars, 20 µm.

**Supplementary Fig. 5.**
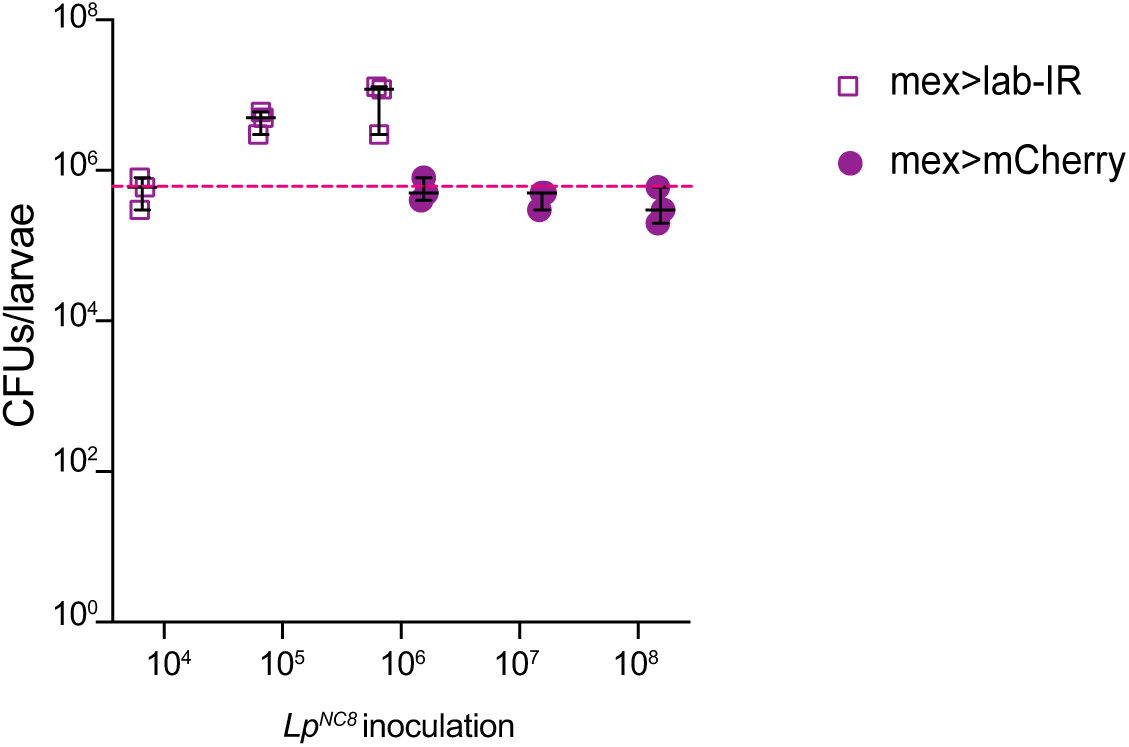
Adjustment of bacterial inoculum to obtain comparable intestinal colonization levels in larvae with or without the acidic gut region. Different initial concentrations of *Lp^NC8^* were inoculated into germ-free *mex>mCherry* controls and *mex>lab-IR* individuals lacking the acidic copper cell region. Bacterial loads (CFUs) were quantified 5 days after inoculation from whole larvae.

**Supplementary Video 1**. Timelapse video of *Lp^NC8^*strain growing on MRS. Cells were placed on agarose pads containing the fluorescent membrane dye FM4-64 (false-colored in magenta).

**Supplementary Video 2.** Timelapse video of *pp2^+^*strain growing on MRS. Cells were placed on agarose pads containing the fluorescent membrane dye FM4-64 (false-colored in magenta).

**Supplementary Video 3.** Timelapse video of *Lp^NC8^*strain growing on MRS. Cells were placed on agarose pads containing the fluorescent membrane dye FM4-64 (false-colored in magenta).

**Supplementary Video 4.** Timelapse video of *Lp^NC8^*strain growing on MRS supplemented with mitomycin C. Cells were placed on agarose pads containing the fluorescent membrane dye FM4-64 (false-colored in magenta).

**Supplementary Video 5.** Timelapse video of *pp2^+^*strain growing on MRS supplemented with mitomycin C. Cells were placed on agarose pads containing the fluorescent membrane dye FM4-64 (false-colored in magenta).

**Supplementary Video 6.** Timelapse video of *Lp^NC8^*_Δpp2lysis_ strain growing on MRS supplemented with mitomycin C. Cells were placed on agarose pads containing the fluorescent membrane dye FM4-64 (false-colored in magenta).

## SUPPLEMENTARY DATA TABLES

**Supplementary Table S1.** Bacterial strains used in this study.

| Strain | Relevant characteristics | Reference or source |
| --- | --- | --- |
| ***E. coli*** |  |  |
| TG1 | *supE hsd5h thi* (Δ*lac-proAB*) *F'* (*traD36 proAB-lacZ*ΔM15) | 56 |
| GM1674 | *dam<sup>-</sup>dcm<sup>-</sup>repA<sup>+</sup>* | 57 |
| *pks<sup>+</sup>* | *E. coli* BW25113 harboring a BAC with *pks* locus | Gift from Eric Oswald |
| *pks<sup>-</sup>* | *E. coli* BW25113 harboring the control BAC construction without *pks* locus | Gift from Eric Oswald |
| ***L. plantarum*** |  |  |
| NC8 | Isolated from grass silage, plasmid free | 58 |
| Δ*recA* | NC8 strain deleted for *nc8_2035* | This study |
| *pp<sup>-</sup>* | NC8 strain deleted for *pp1*, *pp2* and *pp3* | This study |
| *pp1<sup>+</sup>* | NC8 strain deleted for *pp2* and *pp3* | This study |
| *pp2<sup>+</sup>* | NC8 strain deleted for *pp1* and *pp3* | This study |
| *pp3<sup>+</sup>* | NC8 strain deleted for *pp1* and *pp2* | This study |
| *NC8Δpp2lysis* | NC8 strain deleted for *nc8_0567* and *nc8_0568* | This study |
| *pp2<sup>+</sup>Δpp2lysis* | *pp2<sup>+</sup>* strain deleted for *nc8_0567* and *nc8_0568* | This study |
| *pp1<sup>-</sup>* | NC8 strain deleted for *pp1* | This study |
| *pp2<sup>-</sup>* | NC8 strain deleted for *pp2* | This study |
| *pp3<sup>-</sup>* | NC8 strain deleted for *pp3* | This study |

**Supplementary Table S2.**
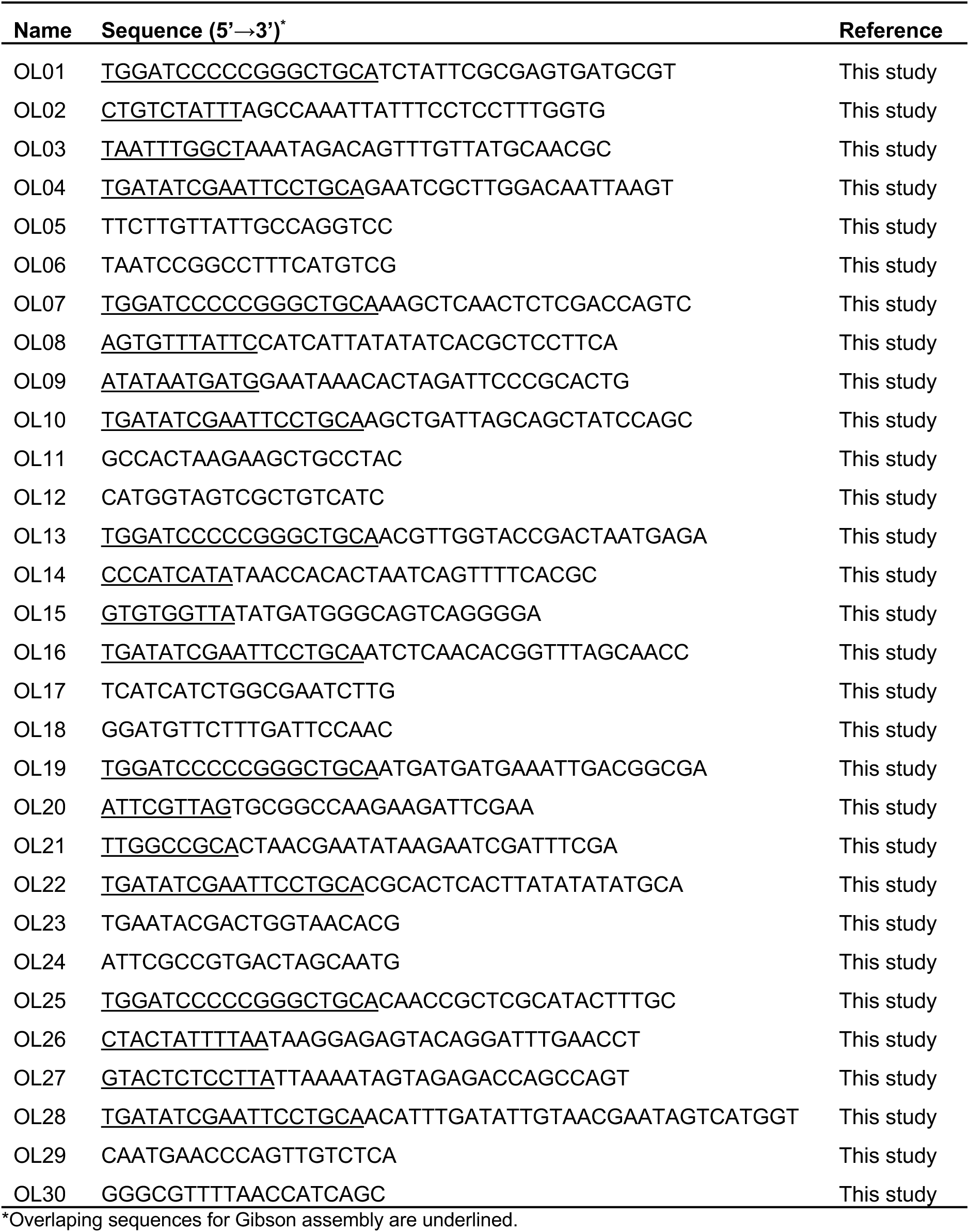
Primers used for the construction of *L. plantarum* strains.

